# From sequence to function through structure: deep learning for protein design

**DOI:** 10.1101/2022.08.31.505981

**Authors:** Noelia Ferruz, Michael Heinzinger, Mehmet Akdel, Alexander Goncearenco, Luca Naef, Christian Dallago

**Affiliations:** Institute of Informatics and Applications, University of Girona, Girona, Spain; Department of Biochemistry, University of Bayreuth, Bayreuth, Germany; Department of Informatics, Bioinformatics & Computational Biology, Technische Universität München, 85748 Garching, Germany; VantAI, 151 W 42nd Street, New York, NY 10036; NVIDIA DE GmbH, Einsteinstraße 172, 81677 München, Germany

**Keywords:** protein design, protein prediction, drug discovery, deep learning, protein language models

## Abstract

The process of designing biomolecules, in particular proteins, is witnessing a rapid change in available tooling and approaches, moving from design through physicochemical force fields, to producing plausible, complex sequences fast via end-to-end differentiable statistical models. To achieve conditional and controllable protein design, researchers at the interface of artificial intelligence and biology leverage advances in natural language processing (NLP) and computer vision techniques, coupled with advances in computing hardware to learn patterns from growing biological databases, curated annotations thereof, or both. Once learned, these patterns can be leveraged to provide novel insights into mechanistic biology and the design of biomolecules. However, navigating and understanding the practical applications for the many recent protein design tools is complex. To facilitate this, we 1) document recent advances in deep learning (DL) assisted protein design from the last three years, 2) present a practical pipeline that allows to go from *de novo*-generated sequences to their predicted properties and web-powered visualization within minutes, and 3) leverage it to suggest a generated protein sequence which might be used to engineer a biosynthetic gene cluster to produce a molecular glue-like compound. Lastly, we discuss challenges and highlight opportunities for the protein design field.

**Availability:** pLM generated and UniRef50 sampled sequence sets and predictions are available at http://data.bioembeddings.com/public/design. Code-base and Notebooks for analysis are available at https://github.com/hefeda/PGP. An online version of **Table 1** can be found at https://github.com/hefeda/design_tools.

## Introduction

Proteins take part in nearly every process of life, controlling a wide variety of functions. This functional versatility and the fact that proteins are nanoscopic, biodegradable materials have motivated tremendous efforts toward designing artificial proteins for medical and industrial applications. In fact, human-designed proteins are widely used across medicine, agriculture and manufacturing, constituting the active form of four out of the top five top selling pharmaceuticals in 2021 [1]. To cite a few more examples, protein engineers have improved the thermostability of a malaria invasion protein for use as a vaccine antigen [2] or developed an enzyme capable of hydrolyzing Polyethylene terephthalate (PET), providing a new, green, and scalable route for plastic waste recycling [3]. Despite the clear objective of designing functional properties, most protein design and engineering strategies have traditionally leveraged a structure-first approach [4], i.e. either by re-engineering known proteins or by designing novel stable structures that then could be further tweaked to target desired functions, e.g. binding to other molecules [5]. The reason why the concomitant design of sequence, structure, and function has remained so challenging is due to the computational intractability of the problem: traditionally, protein design has been tackled as a mathematical optimization, where an algorithm, such as Monte Carlo [6], searched the global minima of a multi-dimensional physicochemical energy function [7]. In recent months, however, an explosion of methods leveraging advances in machine learning provide a fresh alternative for *de-novo* protein design, including the design of long functional proteins. Motivated by the enormous success of structure prediction methods [8]–[11] and the recent availability of large putative protein structure databases [11], [12], some works are exploiting a structure-first approach to designing new folds and proteins [13], [14]. In contrast, others operate *sequence-first*, by training large generative language models on vast sequence databases [15], [16, p. 2], [17, p. 2], [18]. Advancing the field by exploiting different modalities (sequence, structure and even function) is fundamental, as no one modality may be able to explain all cell phenomena necessary for the design of biologics [19].

For comprehensive reviews of protein engineering or machine learning methods for protein research we refer to the works by Yang [20] and Defresne [21]. Yet, with fundamental advances to protein design in a brief period of time, this manuscript attempts to provide an overview of recent work using AI with a focus on the last three years. To showcase the practical uses of the work presented, we engineer a pipeline for the generation of *de novo* protein sequences selectable for tailored properties that may benefit the protein design community, and make use of this novel pipeline to discover sequences that may generate natural products.

In particular, in the following:

1. We describe recent advances in protein design, namely those shifting from a physical-based function paradigm to one that uses end-to-end differentiable architectures for sequence and structure generation. We aim to give practitioners a waymark to novel tools and their intended uses.
2. We offer a novel pipeline capable of generating protein sequences with tailored properties. In short, we couple ProtGPT2 [16], which generates *de novo* sequences, with ProtT5 [22, p. 5], which predicts properties from them in order to discriminate sequences by desired functions.
3. We dig into the pipeline by presenting a use case for the selection of factory proteins with the predicted ability to produce natural products.
4. We discuss challenges to the design of marketable proteins with controllable properties.

### Moving from physicochemical functions to deep neural networks in protein research

The *de novo* design of proteins was traditionally approached as an optimization problem where an algorithm searched the global minima of an energy function [23]. This function would evaluate an astronomical number of sequences for a given backbone, quickly leading to an NP-hard problem that required turning to heuristic algorithms and static pairwise potential energy functions to limit computational complexity [24], [25]. These approaches met enormous success, with a myriad of *de novo* generated proteins in the last 20 years [26]. These designs have remarkably evolved from often short, alpha-helical peptides [27] and bundles [28] to complex multi-domain architectures [29]. Much has been said about physicochemical-based protein design, we refer the readers to these comprehensive reviews [5], [7], [26].

More recently however, deep learning (DL) approaches have provided a new venue for protein design research by showing high accuracy in prediction tasks. Highly publicized progress from one such tool came from DeepMind’s AlphaFold [30] in December 2018 at CASP13 [31], a step-wise pipeline incorporating DL attempted to solve the decade-long problem of protein 3D structure prediction from sequence. The sucessor AlphaFold 2 [9], an end-to-end engineered solution, promoted even more excitement due to its incredible ability to accurately predict protein structures from sequences *in-silico* [32], [33]. End-to-end solutions both in protein predictions and design are of particular interest because they set out to learn complex relationships between a complex output (e.g. 3D structure) and a principled input measurement (e.g. sequence) based on mathematical, often biologically-informed formulation of their relationships. Once trained and validated, these models encode a comprehensive notion of the input and its relationships to the output [34], [35].

In parallel to AlphaFold, natural language processing (NLP) methods were leveraged to learn novel protein representations by learning the protein *language*, offering an alternative route to the *explicit* extraction of evolutionary information from MSAs, historically done by collecting statistics on the co-evolution of residues within MSAs [24]. These models achieved an understanding of proteins by tasking multi-million DL architectures to solve millions of cloze tests from large protein sequence datasets, allowing to encode statistics of protein sequences without supervision on physicochemical or evolutionary relationships [22], [36]. While at first pLM representations (shorthand: *embeddings*) did outperform traditional exploitations of direct physicochemical and evolutionary knowledge [37], [38], quick advancements in better mechanisms to process embeddings led to competitive predictors for protein function and structure [10], [39]–[49]. It is thus becoming clear that pLM embeddings are rich inputs to *downstream* prediction methods of protein function and structure competing with those that exploit MSAs. However, what is encoded in embeddings, including how much *evolutionary information* as defined in MSAs (i.e., residue co-evolution) is implicitly captured, remains a subject of debate, even in light of correlation between MSAs and pLM embeddings in accuracy for 3D structure prediction [40].

While these pLMs excel at embedding representations of input sequences, other types of pLMs (discussed below) are now capable of directly generating new sequences. With increasing experimental validation of “black box” DL methods, the protein design field now has a unique opportunity to generate sequences, interpret DL representations, and build pipelines aiding at various stages of protein design and drug development, from initial library generation to refinement and optimization.

### The deep learning era of protein sequence and structure generation

Previous reviews have focused on describing the types of neural network architectures used in protein design [21], [50]. We focus on the type of problem that these methods attempt to solve: 1) fixed-backbone design (**Fig. 1, Panel 1**), 2) sequence generation (**Fig. 1, Panel 2**), 3) structure generation (**Fig. 1, Panel 3**), and 4) concomitant structure and sequence design most often via *hallucination* (**Fig. 1, Panel 4**). We discuss several DL methods (bold and underlined to match rows **Table 1**) from the last three years with a focus on those not using Potts models [51], [52]. A virtual version of this table can be found at https://github.com/hefeda/design_tools.

**Fig. 1:**
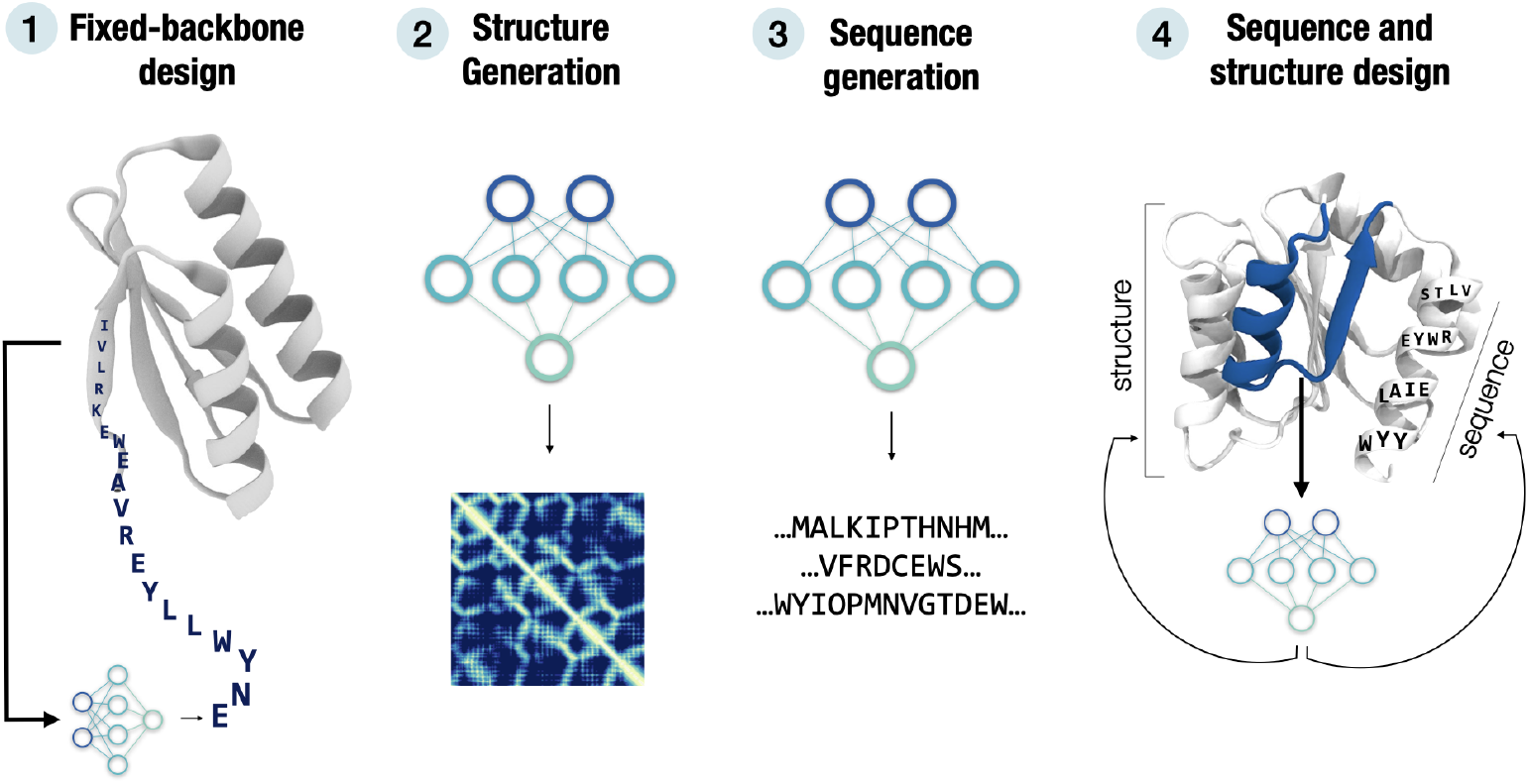
leveraging deep learning for different protein design goals. We classify DL approaches in **(1)** models that try to solve the traditional protein design problem of inverse folding, i.e. find sequences that fold into a desired structure, **(2)** models capable of generating structure-encoding objects, like contact or distance maps, **(3)** models that learn to generate protein sequences or **(4)** concomitantly model sequence and structure to generate either. Categories **1-4** are cross-linked in **Table 1**.

**Table 1:**
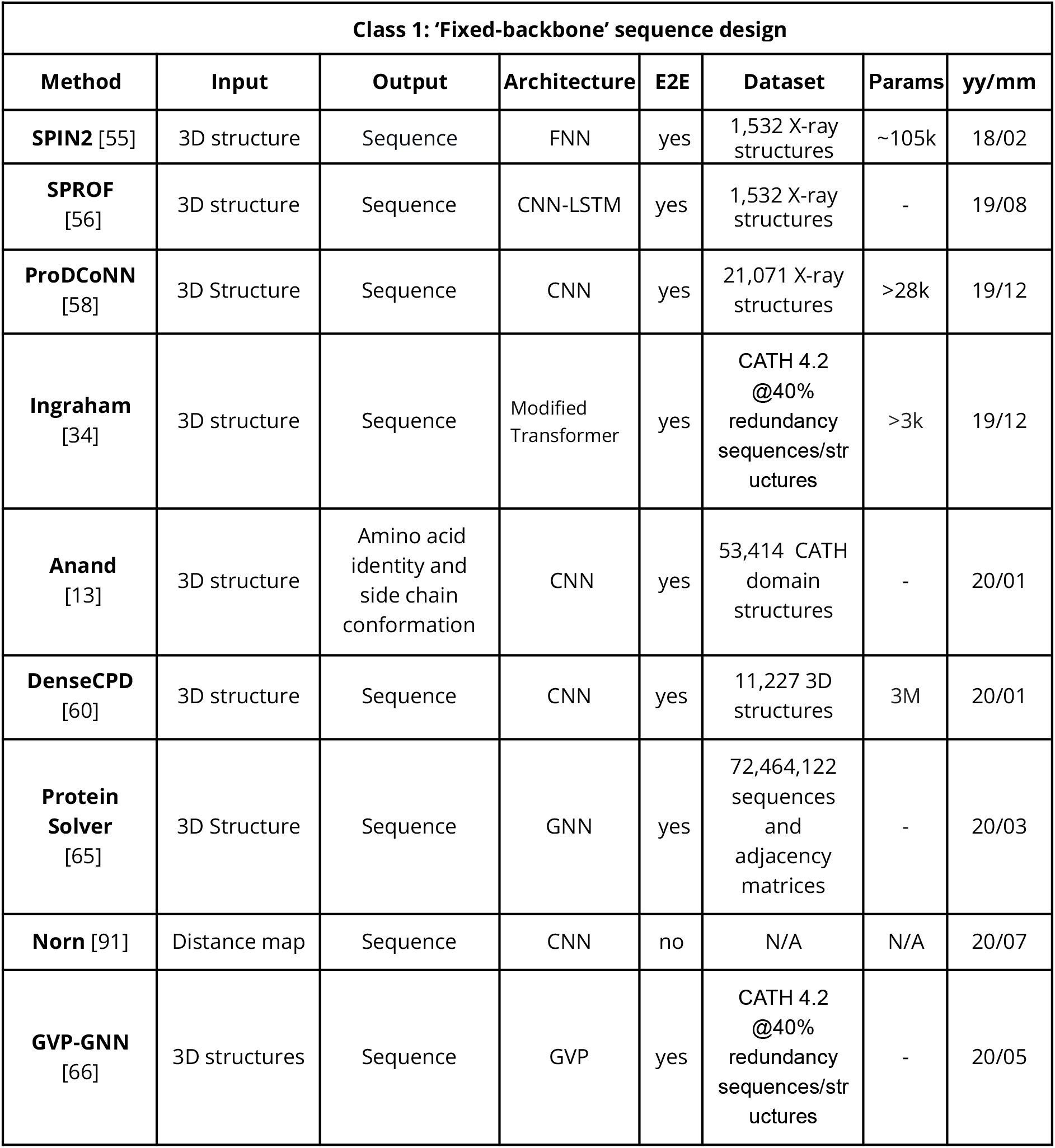
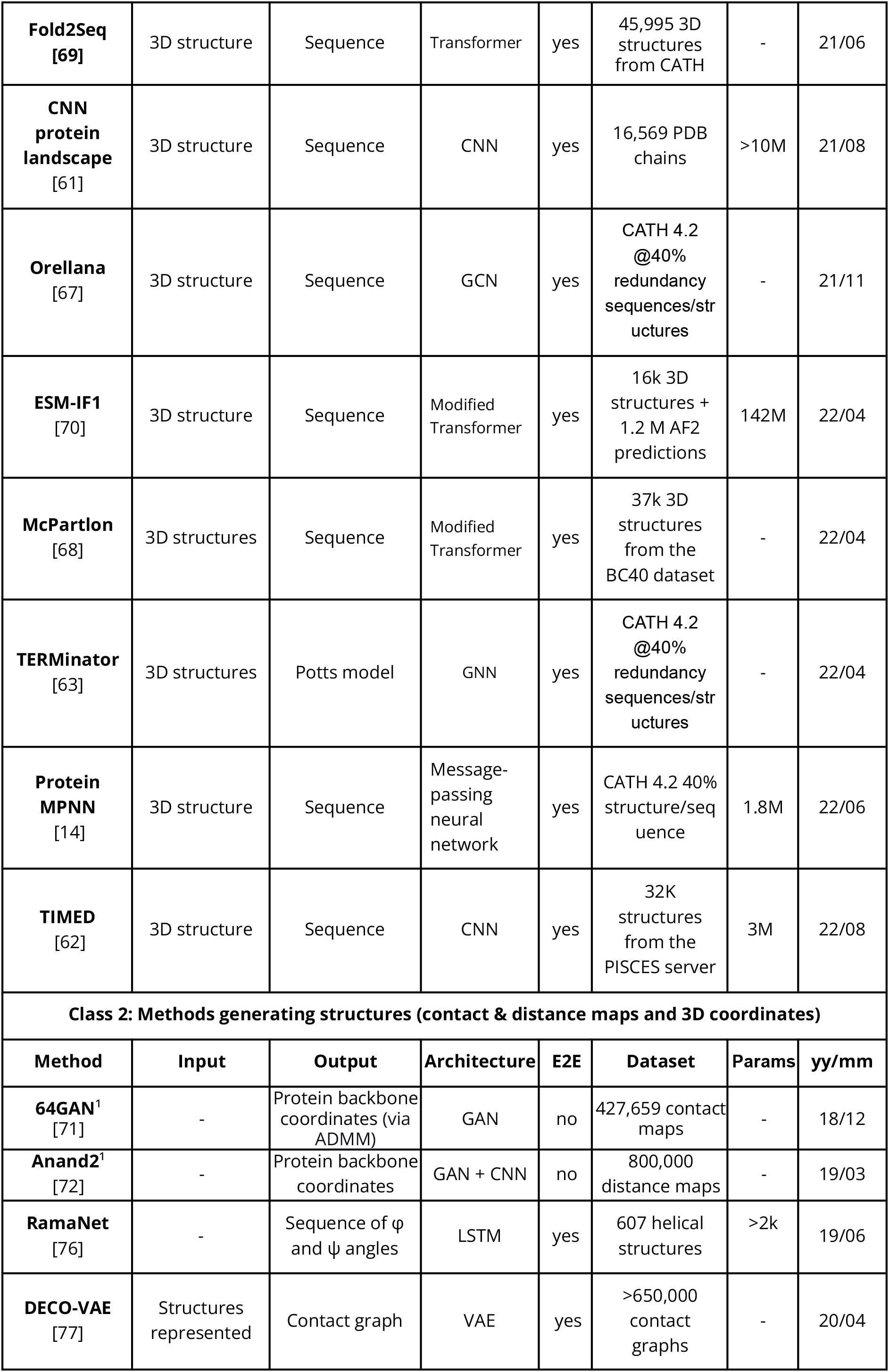
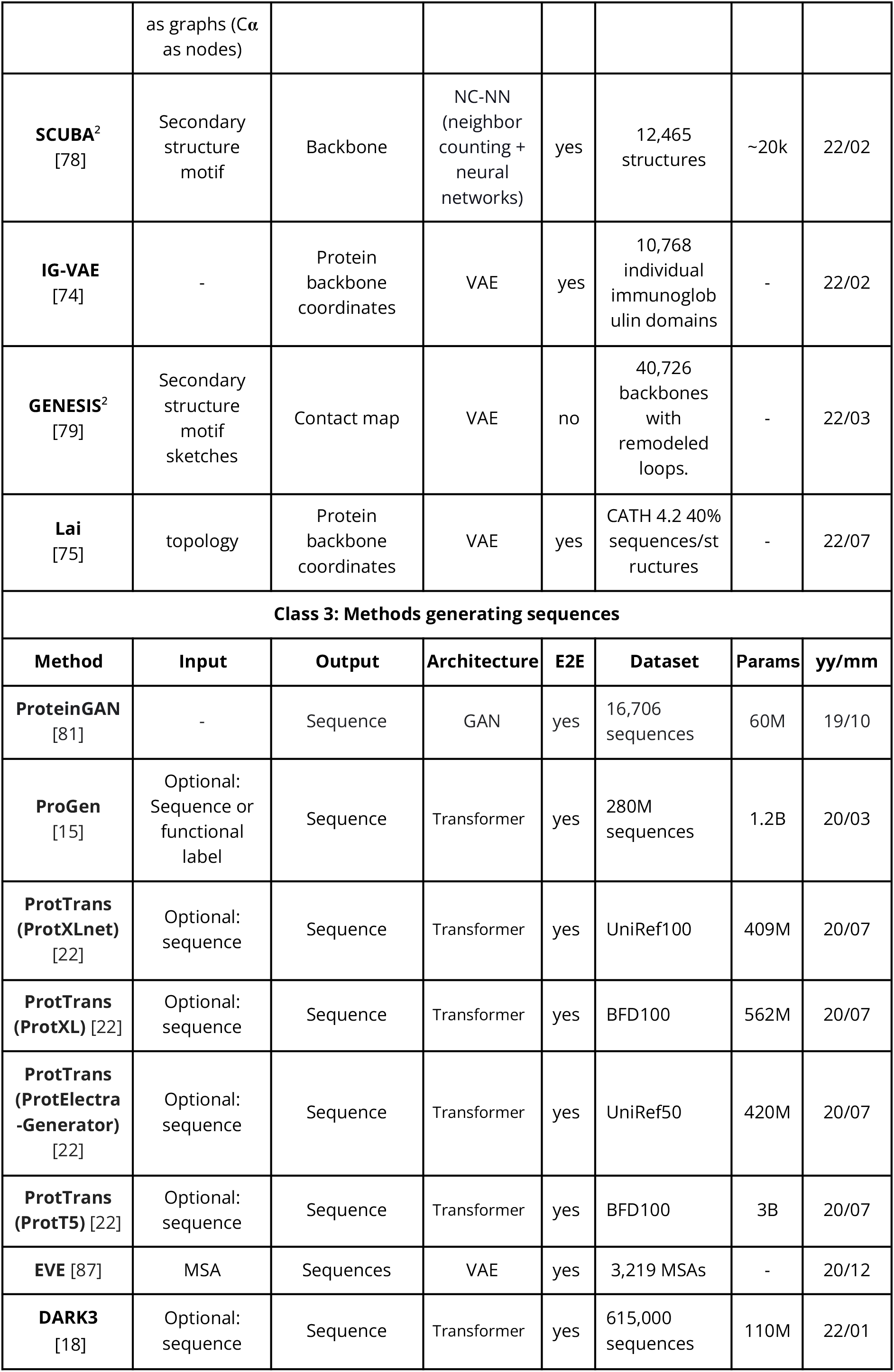
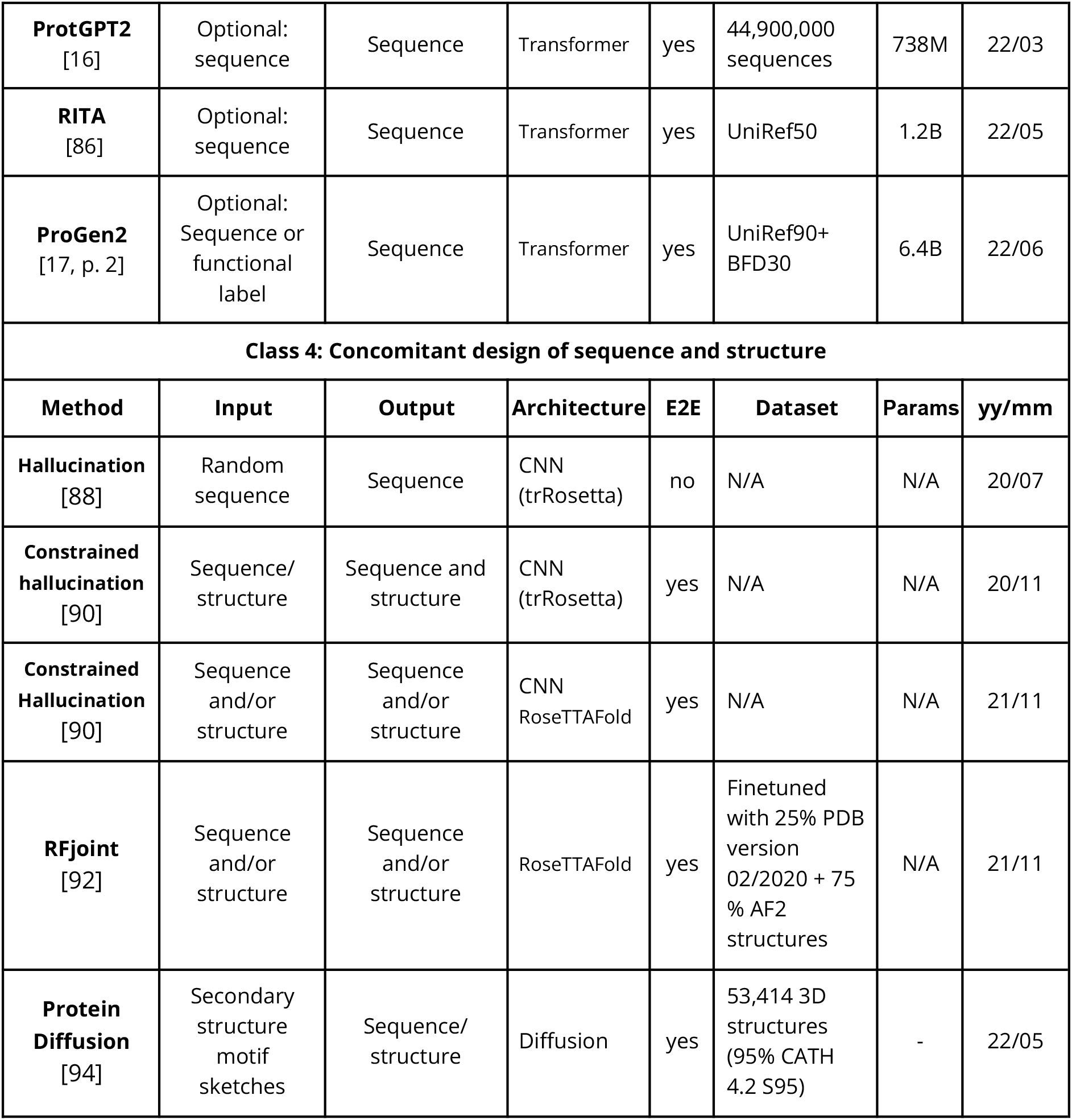
machine-learning-based protein design methods. Methods ordered by their release date, accounting for the date of pre-prints when available. Class (1-4) captured in **Fig. 1**. Expanded method detail in main text. Unnamed methods are referenced by the first name of the first author. Online version at https://github.com/hefeda/design_tools. Legend: **Architecture**: The architecture of the deep learning model; **E2E**: an end-to-end differentiable solution; **Input**: the input to run (infer from) the model, e.g. a contact map; **Output**: the output, e.g. a protein sequence; **Dataset**: the number and type of samples used to train the method; **Params**: the exact or estimated number of parameters of the model; ^**1**^: the 3D recovery was performed in an external, second step; ^**2**^: conditioned generation; –: no input required for generation.

| Class 1: 'Fixed-backbone' sequence design |  |  |  |  |  |  |  |
| --- | --- | --- | --- | --- | --- | --- | --- |
| Method | Input | Output | Architecture | E2E | Dataset | Params | yy/mm |
| **SPIN2** [55] | 3D structure | Sequence | FNN | yes | 1,532 X-ray structures | ~105k | 18/02 |
| **SPROF** [56] | 3D structure | Sequence | CNN-LSTM | yes | 1,532 X-ray structures | - | 19/08 |
| **ProDCoNN** [58] | 3D Structure | Sequence | CNN | yes | 21,071 X-ray structures | >28k | 19/12 |
| **Ingraham** [34] | 3D structure | Sequence | Modified Transformer | yes | CATH 4.2 @40% redundancy sequences/structures | >3k | 19/12 |
| **Anand** [13] | 3D structure | Amino acid identity and side chain conformation | CNN | yes | 53,414 CATH domain structures | - | 20/01 |
| **DenseCPD** [60] | 3D structure | Sequence | CNN | yes | 11,227 3D structures | 3M | 20/01 |
| **Protein Solver** [65] | 3D Structure | Sequence | GNN | yes | 72,464,122 sequences and adjacency matrices | - | 20/03 |
| **Norn** [91] | Distance map | Sequence | CNN | no | N/A | N/A | 20/07 |
| **GVP-GNN** [66] | 3D structures | Sequence | GVP | yes | CATH 4.2 @40% redundancy sequences/structures | - | 20/05 |
| **Fold2Seq**<br>[69] | 3D structure | Sequence | Transformer | yes | 45,995 3D structures from CATH | - | 21/06 |
| **CNN protein landscape**<br>[61] | 3D structure | Sequence | CNN | yes | 16,569 PDB chains | >10M | 21/08 |
| **Orellana**<br>[67] | 3D structure | Sequence | GCN | yes | CATH 4.2 @40% redundancy sequences/structures | - | 21/11 |
| **ESM-IF1**<br>[70] | 3D structure | Sequence | Modified Transformer | yes | 16k 3D structures + 1.2 M AF2 predictions | 142M | 22/04 |
| **McPartlon**<br>[68] | 3D structures | Sequence | Modified Transformer | yes | 37k 3D structures from the BC40 dataset | - | 22/04 |
| **TERMinator**<br>[63] | 3D structures | Potts model | GNN | yes | CATH 4.2 @40% redundancy sequences/structures | - | 22/04 |
| **Protein MPNN**<br>[14] | 3D structure | Sequence | Message-passing neural network | yes | CATH 4.2 40% structure/sequence | 1.8M | 22/06 |
| **TIMED**<br>[62] | 3D structure | Sequence | CNN | yes | 32K structures from the PISCES server | 3M | 22/08 |
| **Class 2: Methods generating structures (contact & distance maps and 3D coordinates)** |  |  |  |  |  |  |  |
| **Method** | **Input** | **Output** | **Architecture** | **E2E** | **Dataset** | **Params** | **yy/mm** |
| **64GAN**<sup>1</sup><br>[71] | - | Protein backbone coordinates (via ADMM) | GAN | no | 427,659 contact maps | - | 18/12 |
| **Anand2**<sup>1</sup><br>[72] | - | Protein backbone coordinates | GAN + CNN | no | 800,000 distance maps | - | 19/03 |
| **RamaNet**<br>[76] | - | Sequence of $\varphi$ and $\psi$ angles | LSTM | yes | 607 helical structures | >2k | 19/06 |
| **DECO-VAE**<br>[77] | Structures represented | Contact graph | VAE | yes | >650,000 contact graphs | - | 20/04 |

From sequence to function through structure: deep learning for protein design.
| | as graphs (Cα as nodes) | | | | | | |
| --- | --- | --- | --- | --- | --- | --- | --- |
| **SCUBA<sup>2</sup>**<br>[78] | Secondary structure motif | Backbone | NC-NN (neighbor counting + neural networks) | yes | 12,465 structures | ~20k | 22/02 |
| **IG-VAE**<br>[74] | - | Protein backbone coordinates | VAE | yes | 10,768 individual immunoglobulin domains | - | 22/02 |
| **GENESIS<sup>2</sup>**<br>[79] | Secondary structure motif sketches | Contact map | VAE | no | 40,726 backbones with remodeled loops. | - | 22/03 |
| **Lai**<br>[75] | topology | Protein backbone coordinates | VAE | yes | CATH 4.2 40% sequences/structures | - | 22/07 |
| **Class 3: Methods generating sequences** |  |  |  |  |  |  |  |
| Method | Input | Output | Architecture | E2E | Dataset | Params | yy/mm |
| **ProteinGAN**<br>[81] | - | Sequence | GAN | yes | 16,706 sequences | 60M | 19/10 |
| **ProGen**<br>[15] | Optional: Sequence or functional label | Sequence | Transformer | yes | 280M sequences | 1.2B | 20/03 |
| **ProtTrans (ProtXLnet)**<br>[22] | Optional: sequence | Sequence | Transformer | yes | UniRef100 | 409M | 20/07 |
| **ProtTrans (ProtXL)** [22] | Optional: sequence | Sequence | Transformer | yes | BFD100 | 562M | 20/07 |
| **ProtTrans (ProtElectra-Generator)**<br>[22] | Optional: sequence | Sequence | Transformer | yes | UniRef50 | 420M | 20/07 |
| **ProtTrans (ProtT5)** [22] | Optional: sequence | Sequence | Transformer | yes | BFD100 | 3B | 20/07 |
| **EVE** [87] | MSA | Sequences | VAE | yes | 3,219 MSAs | - | 20/12 |
| **DARK3**<br>[18] | Optional: sequence | Sequence | Transformer | yes | 615,000 sequences | 110M | 22/01 |
| **ProtGPT2**<br>[16] | Optional:<br>sequence | Sequence | Transformer | yes | 44,900,000<br>sequences | 738M | 22/03 |
| **RITA**<br>[86] | Optional:<br>sequence | Sequence | Transformer | yes | UniRef50 | 1.2B | 22/05 |
| **ProGen2**<br>[17, p. 2] | Optional:<br>Sequence or<br>functional<br>label | Sequence | Transformer | yes | UniRef90+<br>BFD30 | 6.4B | 22/06 |
| **Class 4: Concomitant design of sequence and structure** |  |  |  |  |  |  |  |
| **Method** | **Input** | **Output** | **Architecture** | **E2E** | **Dataset** | **Params** | **yy/mm** |
| **Hallucination**<br>[88] | Random<br>sequence | Sequence | CNN<br>(trRosetta) | no | N/A | N/A | 20/07 |
| **Constrained**<br>**hallucination**<br>[90] | Sequence/<br>structure | Sequence and<br>structure | CNN<br>(trRosetta) | yes | N/A | N/A | 20/11 |
| **Constrained**<br>**Hallucination**<br>[90] | Sequence<br>and/or<br>structure | Sequence<br>and/or<br>structure | CNN<br>RoseTTAFold | yes | N/A | N/A | 21/11 |
| **RFjoint**<br>[92] | Sequence<br>and/or<br>structure | Sequence<br>and/or<br>structure | RoseTTAFold | yes | Finetuned<br>with 25% PDB<br>version<br>02/2020 + 75<br>% AF2<br>structures | N/A | 21/11 |
| **Protein**<br>**Diffusion**<br>[94] | Secondary<br>structure<br>motif<br>sketches | Sequence/<br>structure | Diffusion | yes | 53,414 3D<br>structures<br>(95% CATH<br>4.2 S95) | - | 22/05 |

The first category of methods we describe focus on solving the traditional protein design problem, i.e., finding a sequence that optimally adopts a desired backbone (**Fig. 1, Panel 1**). The performance of these methods is usually evaluated by *native sequence recovery* (NSR), i.e., the percentage of wild-type amino acids recovered for an input sequence by the design method.

While this metric imposes some limitations, given that the identity percentage does not necessarily correlate with expression or functional levels [53] and the lack of common benchmark sequences, it is nevertheless a convenient measure to evaluate how well the method recapitulates wild-type sequences. Some of the first attempts came from SPIN [54] and SPIN2 [55, p. 2], which leveraged three-layered fully-connected neural networks (FNN) to learn from structural features embedded as a 1-dimensional (1D) tensor representing backbone torsion angles, local fragment-derived profiles, and global energy-based features. While SPIN and SPIN2 achieved an NSR of 30 % and 34%, respectively, they suffered from information loss due to the 1D input representation and the lack of FNNs to encode local and global context. Their successor, SPROF [56], tried to remedy information loss by leveraging a 2D input and a 2D Convolutional Neural Network (CNN), which brought NSR to 39.8%. 2D CNN models excelled for years in computer vision tasks as they handle local and global information better than FNNs [57]. In the protein context, these may be spatial and geometric features of 3D structure, which can be extracted in an unsupervised fashion through CNNs, given that protein 3D structures can be proxied accurately (up to symmetry) by 2D representations like contact- or distance maps. For instance, ProDCoNN [58] learned to reconstruct sequences by representing atomic discretized environments through voxelized 18 Å-cubic boxes, leading to a NSR of over 45%. Anand [13]‘s tool designed sidechain rotamers besides recovering sequence identity at a given input backbone position, reaching an NSR of 57%. Three of the designs were validated with X-ray crystallography. Meanwhile, Deep Learning researchers attempted to solve training challenges of CNNs, such as the elevated computational cost associated with increasing neural network depth. For instance, DenseNet [59] expanded on the concept of residual connections, i.e. wiring a layer’s input to all subsequent *residual blocks*, which inspired DenseCPD [60]. Other attempts tried to encode more information in the input to CNNs, e.g. CNN protein landscape [61], which amongst others encoded side chain atoms, partial charges and solvent accessibility reaching 60% NSR, and TIMED [62], which also includes reimplementations of several CNN-based protein design methods.

Protein 3D structures can also be represented by graphs where nodes are residues (or atoms) and edges represent structural proximity. Graph neural networks (GNN) directly harness graph representations, and thus attempts like TERMinator [63], [64] or ProteinSolver [65], a GNN inspired by Sudoku problems, emerged. ProteinSolver generated sequences from four different folds, two of which were experimentally characterized with circular dichroism. Ingraham [34] trained a encoder-decoder *Structured Transformer*, where the GNN encoder learnt protein structures represented by graphs, while the decoder sampled sequences conditioned on the encoder-learn structure representations. Graph Vector Perceptrons (GVP-GNN) [66] which replaced MLPs in GNNs improved performance in protein design and model quality assessment. Through modifications of the GVPs architecture, Orellana [67] improved the median sequence recovery from 40.2% to 44.7%. In recent work, McPartlon [68] introduced a partial masking scheme and side-chain conformation loss to GNNs achieving an NSR of 50.5% on independent CASP, CATH and TS50 test sets. Other recent studies leveraged encoder-decoder Transformer architectures, e.g. Fold2seq [69] learned protein representations jointly from sequence and structure through two encoder modules connected to a decoder module tasked with sampling sequences from the learnt sequence and structure representations. ESM-1F [70] used GVP-GNN encoder layers to represent geometric features, followed by a generic autoregressive encoder-decoder, improving performance over previous GVP-GNN networks. ProteinMPNN [14] implemented a Transformer whose encoder embeds the protein backbone coordinates, whilst the decoder outputs suitable sequences. The method was experimentally validated, showing high expression yields (in some cases rates over 88%) across different tasks. One design was crystallized, showing a more complex fold than most *de novo* proteins up to date [14].

Most of the early work in the second category of methods generating structure encoding objects (**Fig. 1, Panel 2**) came from the Po-Ssu Huang lab, which initially trained generativ adversarial networks (GANs) on contact maps used as input to a convex optimization algorithm (ADMM) to recover 3D coordinates [71]. Later, the network was improved in Anand2 to learn from distance maps, and included a learned coordinate recovery module replacing ADMM [72]. In these methods, sequences were generated from the designed backbones using Rosetta [73]. GANs were subject to a lack of satisfactory accuracy in the coordinate recovery process and the loss of resolution when inputting structures as either contact or distance maps, which led to missing biochemical features and unrealistic designs [74]. IG-VAE [74] addressed some of these shortcomings by training a variational autoencoder (VAE) that directly generated 3D coordinates of backbone atoms for class-specific Immunoglobulin proteins. The VAE implemented in Lai [75] instead outputs a conformational ensemble of protein structures. Similar methods with the goal of generating structures are RamaNet [76], a long-short term memory network (LSTM), trained in an autoregressive manner to output a sequence of φ and Ψ angles to design alpha-helical structures, and DECO-VAE [77], based on VAEs. As opposed to the fixed-backbone methods (**Fig. 1, Panel 1**), these generative structure methods allowed to explore novel, unseen topologies, which could host novel functions. Nevertheless, it is often desirable to control the design process, i.e., to condition the sampling towards aspects of function, structure, or sequence. Two methods (SCUBA and GENESIS) allow to do so through inputting a series of secondary structure rules of sketches (such as ‘helix-loop-helix’), which guide the network. SCUBA [78] used statistical representations of backbones by tasking FNNs to learn from radial kernels encoding different representations of 3D structure. Designs from SCUBA were experimentally evaluated through X-ray crystallography, leading to the discovery of three novel topologies. GENESIS [79] implemented a VAE that takes secondary structure sketches and outputs contact maps with finer definitions of secondary structural elements.

Another emerging protein design branch focuses on sequence generation (**Fig. 1, Panel 3**), mostly inspired by the impressive advances in natural language processing (NLP) over the last few years [50], [80]. Possibly the first advance in this area came from ProteinGAN [81], a GAN trained on the family of malate dehydrogenases (MDH) capable of producing novel functional MDH sequences with as low as 66% identity to natural protein sequences. Since then, a myriad of autoregressive protein language models (pLMs) with generative capabilities, often leveraging the successful Transformer architecture [82], have followed. Since the applications of autoregressive Transformers to create protein language models have extensively been reviewed [50], we will only briefly mention these. ProGen [15] was the first reported decoder-only model specifically trained for protein sequence design and included over 1,100 UniProt [83] control tags (*keywords*). These tags could be used to control the generation process, e.g. by selecting for acyltransferase activity. Transformer models can also be “*fine-tuned*” to achieve a desired goal, practically an alternative technique to using tags for controllable generation. ProGen was employed for the generation of Lysozimes using fine-tuning on five fold diverse Lysozimes families resulting in about 50.000 sequences. Generated sequences showed enzymatic activity in the range of natural counterparts, and one sequence was purified and its structure resolved via X-ray [84]. ProtTrans [22], an extensive probe into the ability of six transformer-based architectures to encode protein sequence knowledge, included the training of autoregressive models (ProtXL, ProtXLNet, ProtElectra-Generator-BFD, and ProtT5) which have, in principle, sequence generation ability. DARK3 [18] is a decoder-only model with 100M parameters trained on synthetic sequences. Following the principles of DARK3, ProtGPT2 leveraged a GPT2-like model [85] an trained on the Uniref50 dataset [83], leading to a model able to generate proteins in unexplored regions of the natural protein space, while presenting natural-like properties [16]. RITA [86] included a study on the scalability of generative Transformer models with several model-specific (e.g. perplexity) and application-specific (e.g. sequence properties) benchmarks, similar to ProGen2 [17, p. 2], which was accompanied by the release of all pre-trained weights and architectures, ranging from 151M to 6.4B parameter models. Another model with generative ability is EVE [87], a VAE used to predict the pathogenicity of protein variants.

The last category of design methods encompasses those capable of concomitantly designing sequence and structure (**Fig. 1, Panel 4**). Possibly the first method in this class, Hallucination [88] allowed to “*hallucinate”* 100-amino acid long de-novo protein sequences leveraging trRosetta. The term *hallucination* was inspired by DeepDream [89], a CNN capable of generating mesmerizing, psychedelic images by combining input patterns iteratively (e.g. generating a house from eyes and faces). Protein hallucination works similarly: random sequences (which only have arbitrary local structural patterns) are passed to a structure prediction method, such as trRosetta, which predicts a distance map. The difference between this map and a background distribution trained on high-resolution natural structures is iteratively minimized by mutating the sequence one mutation at the time and re-computing its distance map in order to ultimately reach a minimal (or optimal) distance between background distribution and distance map. Iterating over this process for 40,000 steps using Monte Carlo led to sharply defined distance maps. 129 designs were expressed in E.coli, of which 27 were monomeric and well-folded, with three being validated through crystallization [88]. Similar to other applications, it is often paramount to gain control over specific properties, such as building a scaffold around a particular structural motif, like a binding pocket. Constrained hallucination [90] modified the hallucination process in two ways: first by using a composite loss function, combining the losses from Norn [91] and Anishchenko [88], which allowed to create a model to concomitantly find a sequence for the structural motif, while hallucinating the scaffold around it; second, the Monte Carlo sampling procedure was replaced by a gradient-based sequence optimization, leading to an 18-fold decrease in sampling time (from 90 to 5 minutes). In RFjoint [92], this approach was further enhanced by employing RoseTTAFold [93] instead of trRosetta, thus leading to finding a minimum for 3D structure differences instead of distance maps [92]. However, this constrained hallucination schema was still too computationally expensive, and thus RFjoint also fine-tuned a RoseTTAFold variant with a three-term loss which allowed inpainting missing sequences and structures in a few seconds. The method was extensively evaluated experimentally [92]. Lastly, Protein Diffusion [94] leveraged diffusion models [95]–[97] popularized by generative image approaches [98] to train a model capable of generating sequence and structure based on a set of secondary structure input constraints.

### An offline pipeline for protein generation and selection through visual exploration of predicted features

Despite significant advances, the field appears to remain distant from tools allowing the seamless design of proteins that fulfill specific properties in an end-to-end fashion, and even farther for *the end* to be marketable devices primed to ace clinical trials or pass environmental regulations. A step towards this goal would be DL models that prompted with a set of biological mechanism and industry-relevant properties (e.g., desired thermostability, aggregate viscosity, sequence length, subcellular localization, catalytic capabilities, or binding partners) output a sequence or structure satisfying the selected criteria with high precision in a timely fashion. Whilst this may not yet be possible, we present an offline attempt (i.e., not an end-to-end solution) by combining a generative DL model producing sequences [16] with an oracle discriminator DL model helping to query generated sequences for desired properties [22].

In particular, we unconditionally (i.e., without priors on family, function or structure) generated a set of 100,000 protein sequences using ProtGPT2, and predicted secondary structure [22], Gene Ontology (GO) terms [44], residue ability to bind small molecules, nucleotides or metals [47], protein subcellular localization [46], transmembrane topology [41], residue conservation [42], residue disorder [43] and CATH family [45]. Remarkably, this generated a repertoire of 100,000 protein sequences with ∼12 predicted features of structure and function from a single script in approximately 3.5 hours (**Supplement 1**). To analyze whether the generated sequences resemble natural ones, we prepared two more sets to compare against: 1) 100,000 sampled protein sequences from UniRef50 [83] which we call “*U50”* and 2) a *nonsens*e sequence set created by shuffling residues within the sequences of the sampled UniRef50 set (e.g. “SEQVENCE” may become “QEEVNCSE”) which we call “*random”*. We then predict structure and function using the same methods and script applied to the generated sequences for both sets.

Qualitatively comparing distributions of several predicted features for the three sets (**Fig. 2**) suggests that generated protein sequences resemble natural sequences more than random sequences. As generated sequences are accompanied by a wealth of interpretable predicted features, we provide a simple Jupyter Notebook (https://github.com/hefeda/PGP) that allows to query the generated protein set in *quasi-*natural language via the predicted features. To cite two examples, a user could shortlist protein sequences with more than 200 residues that have at least 30% of residues involved in alpha-helical secondary structure and that locate on the outer cell membrane according to GO annotations, or select for sequences shorter than 100 residues with transmembrane strand content and binding to small molecules. The resulting sequences filtered by the desired query displayed in the Jupyter Notebook are linked to LambdaPP [99], a web-server for visual exploration of predicted features that also predicts and displays protein 3D structure using ColabFold [100] and allows 3D structure comparisons through FoldSeek [101].

**Fig. 2:**
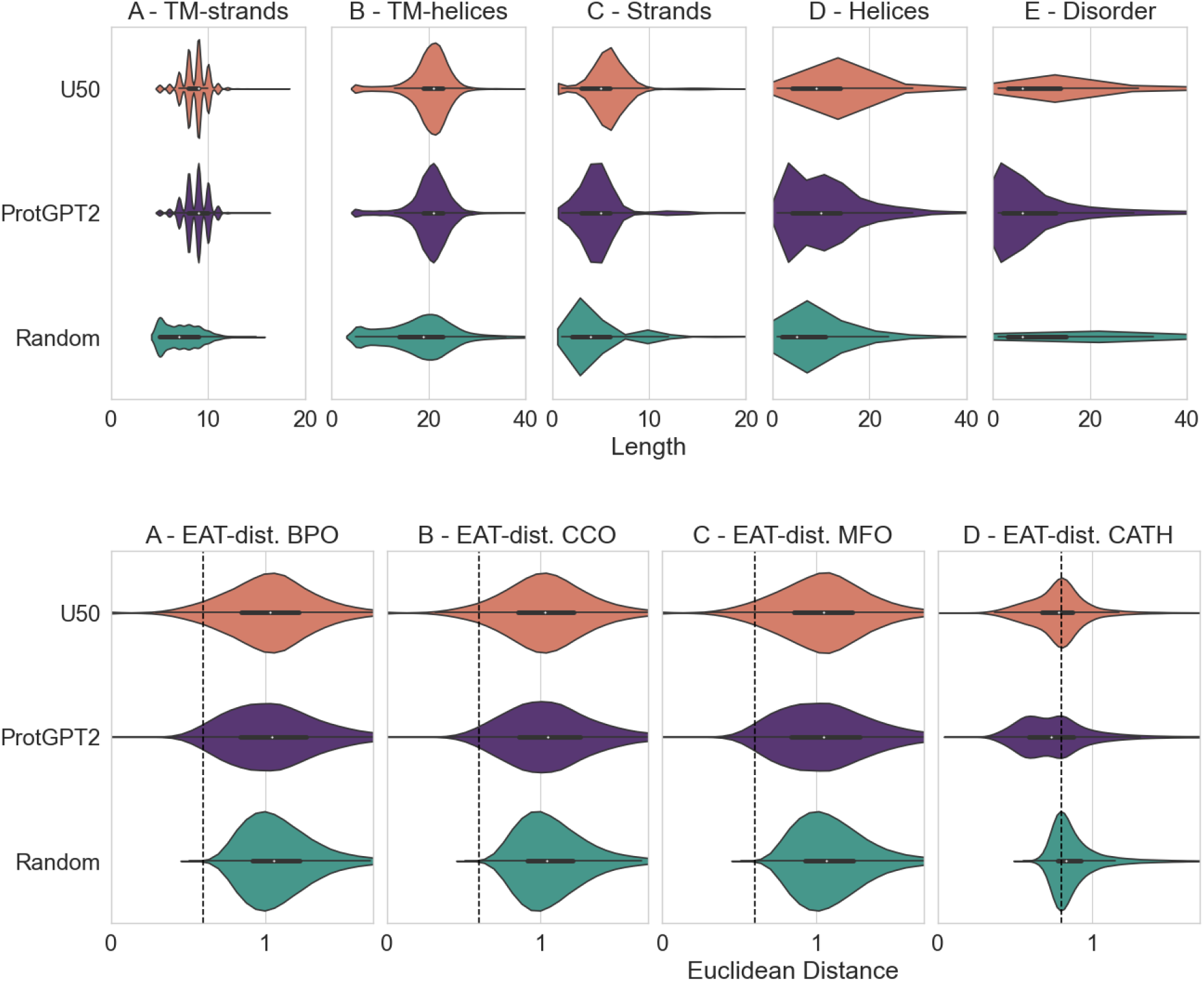
distributions of predicted structural and functional features suggest generated proteins closer to natural sequences than random. Legend: **U50/**orange: 100k sequences sampled from UniRef50; **ProtGPT2/**violet: 100k Dl-generated sequences; **Random/**green: sequences created by randomly shuffling the residues within the U50 set; **Length**: length of the stretch, e.g. 10 consecutive residues; **Euclidean Distance**: the Euclidean distance in embedding space to the first sequence in an annotated set. Structure at residue resolution: consecutive residues forming stretches falling into predicted transmembrane strands (**A**), helices (**B**), or general strands (**C**), helices (**D**) or disorder (**E**) follow similar stretch length distribution between U50 (orange) and ProtGPT2 (violet), but differ from random (green). For instance, random sequences have a preference for short stretches (**B-E**, left of distribution) and are much less likely to be predicted as part of a TM-strand or -helix (**A-B**). Function at sequence resolution: embedding-based annotation transfer (EAT) distribution of distance between proteins in U50, ProtGPT2 and Random to the closest protein with existing GO annotations taken from SwissProt of either biological processes (BPO/**F**), cellular compartment (CCO/**G**) or molecular function (MFO/**H**), or to sequences with CATH annotations (**I**). The distance distribution in the high confidence interval (up to euclidean distance of 0.6, indicated by black, dashed line) is similar for U50 (orange) and ProtGPT2 (violet), but differs for random (green). Comparing the three sets suggests that annotation transfers at euclidean distance greater than 0.6 in embedding space (right of vertical dashed line, **F-I**) invite uncertainty, whilst transfers between proteins at euclidean embedding distance lower than 0.6 may be considered reliable.

Whilst not an end-to-end solution, carefully crafted combinations of lightning-fast DL generators and predictors, easy query mechanisms, and rich visual tools allow to push the envelope in discovery of novel candidates, as explored in the following use case.

### Use case: selecting protein factories for molecular glues through deep learning

Proteins can serve as factories increasingly engineered to manufacture other products, such as the triterpene squalene [102]. Often protein factories are produced by genes in biosynthetic gene clusters (BGC) found in many microorganisms. Protein language model-based deep learning was used to create a tool able to scan genomes for BGCs, assess their functional classification according to Pfam [103], and predict their output product [104]. These protein factories are also the source of many therapeutically used natural products, of which an estimated 50% of all FDA approved drugs are derived [105]. It was discovered that many of these natural products act through a newly established mechanism [106] used by microbial, plant, fungal and animal cells to regulate protein interaction networks at the proteome scale [106], such as plant hormones Auxin and Jasmonate [107]. These small molecules are now aptly named *molecular glues* as they work by gluing a protein of interest to a regulating protein, often an enzyme called the *effector* protein. Multiple approved drugs with previously unknown mechanisms of action have now been identified to work through this mechanism [108], [109]. Molecular glues hold tremendous promise, allowing to drug protein classes that are traditionally considered undruggable, as they don’t require binding enzymatically active sites, can work in shallow, featureless pockets [110], as well as on intrinsically disordered proteins which can become ordered at the stabilized protein interface [111], [112]. Approved molecular glues, however, have all been discovered through serendipity, and it appears that nature has been much more successful in designing this class systematically.

To show the therapeutic promise of artificially designed proteins, we mine unconditionally ProtGPT2-generated protein sequences for their potential to act as factories to produce novel molecular glues. These sequences could be used as a starting pool in a directed evolution approach. Generating sequences, in contrast to the typically used point mutation and shuffling strategies, can achieve a better balance between exploration of sequence diversity and maintaining reaction stability. To do so, we filtered the set of ProtGPT2-generated sequences discussed previously using embedding-based annotation transfer (EAT) of Gene Ontology (GO) annotations [44] to filter 1,345 sequences matching either the GO term “*secondary metabolite biosynthetic process*” (GO:0044550) or its children terms (including those sequences that match up to embedding distance < 0.6). Using a reference embedding set of sequences from the annotated biosynthetic gene cluster dataset (MiBIG) [113], we discover that even with no further optimization, 234 out of the 1,345 generated, filtered sequences are similar to naturally occurring BGC proteins (**Fig. 3**).

**Fig. 3:**
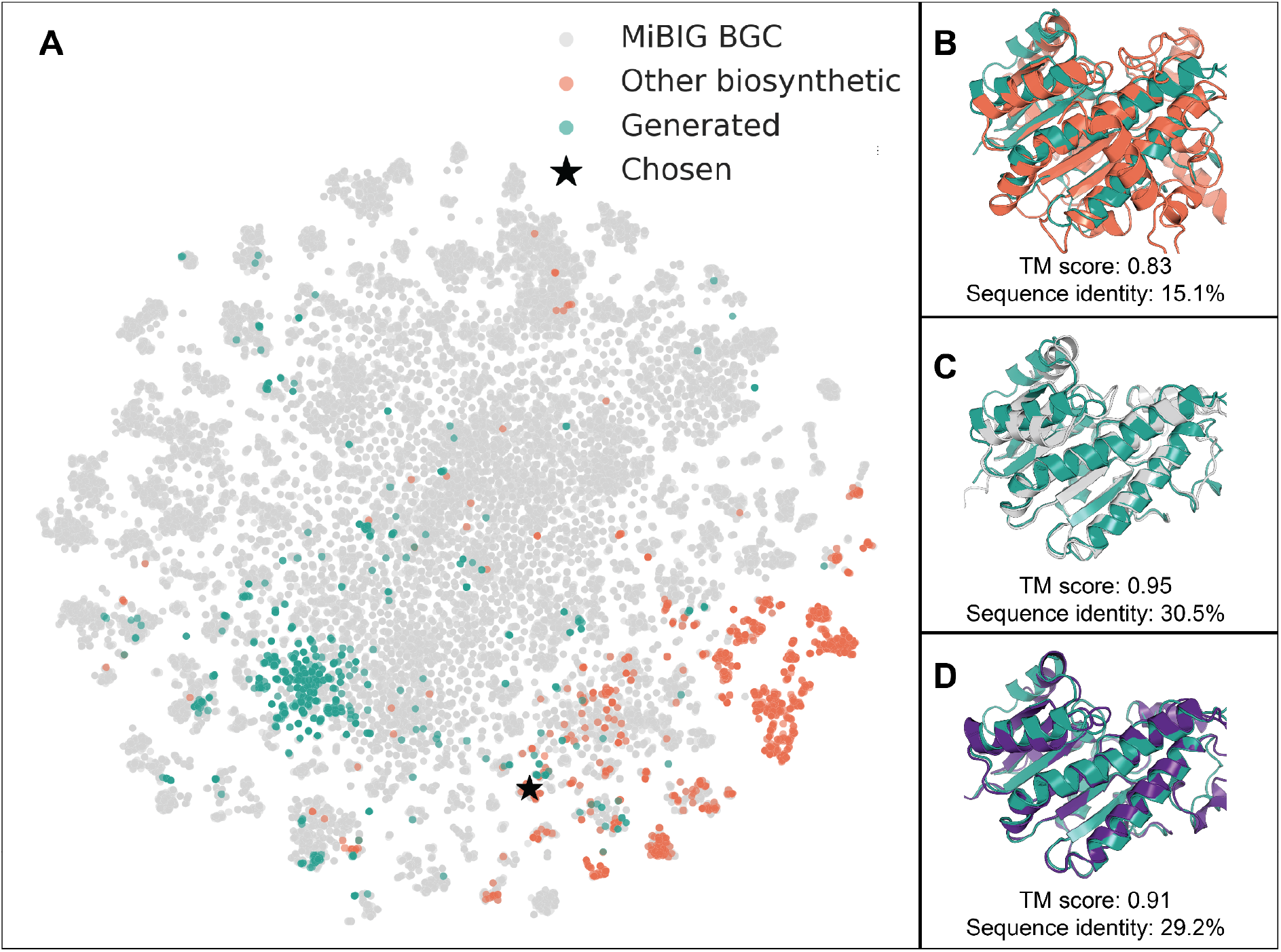
unconditionally generated protein sequences close to natural sequences with desired function. t-SNE projection of protein embeddings for enzymes in biosynthetic gene clusters (as taken from MiBIG [113]; gray), other biosynthetic enzymes (from UniProt; orange), and generated sequences in green (panel A). On the left the structural superposition of the chosen protein (★ in panel A and green in B-D) with the closest biosynthetic protein from UniProt (accession: D7UTD0; orange; panel B), the closest BGC protein from MiBig (UniProt: D2IKP3, MiBIG: BGC0000304|9); gray; panel C), and the closest protein structure from the AlphaFold database [12] found via FoldSeek [101] (id: AF-Q00674; purple; panel D).

We focus on one generated sequence (sequence in **Supplement 1**) which is close to a short chain dehydrogenase BGC constituent involved in producing fusicoccin A (one of the molecules depicted in **Supplement 2, Fig. S1, panel A**). Fusicoccin A is a phytotoxic molecular glue which was originally discovered to stabilize the interface between the scaffolding protein 14-3-3 and H+-ATPases from Arabidopsis thaliana (AHA2) and Nicotiana plumbaginifolia (PMA2) [114], [115], but has since then been expanded semi-synthetically as an anti-cancer drug [116]. By searching for similar proteins in embedding space (**Fig. 3, panel B and C**) and fold space (**Fig. 3, panel D**), three natural proteins emerged. All three are fungal, short-chain dehydrogenases with very similar folds, but are involved in the production of completely different natural products, as expected from different BGCs. This demonstrates a potential workflow of generating sequences with the aim of evolving specific BGCs to produce desired molecules, for instance by applying directed evolution [117].

## Discussion

### Deep learning ushers in a new wave of tools for protein design, engineering, prediction, and optimization

The efficient combination of *in-silico* and *in-vitro* approaches in multi-stage pipelines, such as in drug discovery, remains complex, costly and often requires many types of expertise. However, practitioners now have access to an increasing collection of *in-silico* DL tools which particularly address the first stages of these pipelines (**Table 1**). DL methods have started a revolution in protein design, a field that has undergone a paradigm shift over the last few years, moving away from traditional physico-chemical energy functions. As we highlight in the use case, generative sequence models can be employed to create large libraries of plausible sequences for which oracle models predict structure and function properties in a matter of hours (**Fig. 2**). This process requires no technical DL expertise and provides data from which experts can select and refine to one or a set of promising candidates that constitute starting points for *in-vitro* experiments (**Fig. 3**).

### End-to-end protein design for marketable biologics remains remarkably complex

Despite advances in end-to-end DL models for protein design, except for a few recent breakthroughs [118], bridging the gap from *computational* to *marketable* remains difficult, an unfortunate reality since proteins would have the potential to tackle many emerging biomedical and environmental challenges [119].

Two interesting aspects of DL solutions are their *end-to-end* nature and the ability to combine different losses, i.e. to condition the output (e.g. sequence) on the input (e.g. structure) using a mathematical formulation which takes into account aspects (e.g. thermostability) interesting in the application context (e.g. biopharma). DL models for protein design are optimized end-to-end on large biological collections, which are often limited by experimental shortcomings, for instance capturing a static notion of protein 3D structure, as opposed to its dynamic nature (e.g., conformational modifications during protein-protein interactions). Thus, while DL models for protein design may encode biological mechanisms required to propose viable biologics, they cannot yet be blindly utilized without experimental validation, and do not yet explicitly account for marketable variables. Similarly, as protein function lacks rigorous definition (e.g., may be binding or localization, or both) and scale [120], designing around protein function remains more challenging than designing around sequence or structure, and selecting for function in production is often better validated through targeted functional assays.

An opportunity to connect *in-silico* with *in-vitro* approaches, where physical experiments, accounting for all variables, inform DL models in a differentiable manner may come from advancements in automated labs [121], [122] and a new frontier of cloud labs [123]. However, a bottleneck from combining DL models with physical experimentation is high-throughput [120], as digital systems often vastly outpace physical counterparts in efficiency and scale. Yet, digital copies of physical labs (or digital twins [124]), which have already been proven useful in boosting manufacturing [125], [126] provide an opportunity for the future. A DL solution aiming at outputting marketable devices should also account for non-biological variables, such as the ones influencing production costs or clinical response. Data necessary to inform such models is however lacking, although improvements are underway, such as for clinical trial design [127].

### Looking forward: protein design community on a rigorous journey to establish goals

In the first half of the 1990s, at a time when having *solved the protein folding challenge* sporadically made headlines, the Critical Assessment of Structure Prediction (CASP) [31] set the standard against which *in-silico* predictive methods for protein structure needed to prove advancement. In combination with establishing data standardization and a single data repository, this led to multiple revolutionary approaches, from early uses of single-frequency models (i.e., positional scoring matrices), to complex co-evolutionary representations through direct-coupling analysis [24], all the way to end-to-end DL solutions like AlphaFold 2 [9]. Arguably, the success of structure prediction is a combination of many factors, including technical, biological understanding and intuition, and more complex and principled statistical methods. Fundamentally, however, structure prediction through CASP sets an example of how innovation can be fostered.

As the protein design field moves to more complex DL approaches, benchmarks, which promoted the success of structure prediction tools, appear to be lacking. Arguably, this is due to the underlying complexity of defining, in principle, what protein design is supposed to be, especially as it is moving from theoretical exercise to practical applications. Is it sequence design? Is it structure design? Is it sequence and structure that culminate in function? In fact, function would most likely be the design goal from a practical standpoint. Attempts at measuring advancements in protein function prediction exist, e.g. CAFA [128] and CAGI [129], however they focus on scoring how *in-silico* tools predict known function from sequence, rather than their ability to infer proteins (sequences or structures) that perform a desired, sometimes non-naturally found function. Conversely, model developers score their tools by metrics like Natural Sequence Recovery (NSR), which validate a model’s ability to link structure to sequence, but often not a model’s ability to generate diverse sequences fitting a desired structure [53]. A recent push in benchmarks scoring models’ ability to engineer proteins [130]–[134] highlights three aspects that protein design tools should strive to solve: 1) emulate laboratory conditions, i.e., extrapolate from very little available data; 2) set multiple function generalization goals, i.e, measure different aspects of function, with the intent of finding an optimal solution rather than maximizing any one metric; 3) focus on the ability to address out of observed data distributions, i.e., design proteins that achieve functions not observed in nature.

Ultimately, by overcoming the challenges of measuring advances, deep learning is set to enable protein engineers to design sequence, structure, and function with controllable properties.

## Abbreviations

ADMM: Alternating Direction Method of Multipliers
CNN: Convolutional Neural Network
DL: Deep learning
FNN: fully-connected neural network
GAN: Generative Adversarial Network
GCN: Graph Convolutional Network
GNN: Graph Neural Network
GO: Gene Ontology
GVP: Geometric Vector Perceptron
LSTM: Long-Short Term Memory
MLP: Multilayer Perceptron
MSA: Multiple Sequence Alignment
NLP: Natural Language Processing
NSR: Natural Sequence Recovery
pLM: protein Language Model
VAE: Variational Autoencoder.

## Acknowledgements

We wish to thank Burkhard Rost for inspiring conversations and hardware support. We extend gratitude to all who deposit experimental data in public databases, to those who maintain these databases, and those who make analytical and predictive methods freely available. Additionally, thanks to Simon Duerr, Gina El Nesr, Sergey Ovchinnikov, Kevin K. Yang and all those curating scientific knowledge into easy to parse lists. NF acknowledges support from a Beatriu de Pinos MSCA-COFUND Fellowship (project 2020-BP-00130). CD acknowledges support by the Bundesministerium für Bildung und Forschung (BMBF) through the program “Software Campus 2.0 (TU München)” – project number 01IS17049, and the library of the Technical University of Munich for support in open access costs.

## References

[1] B. Buntz, “50 of 2021’s best-selling pharmaceuticals,” Drug Discovery and Development, Mar. 29, 2022. https://www.drugdiscoverytrends.com/50-of-2021s-best-selling-pharmaceuticals/ (accessed Aug. 15, 2022).

[2] I. Campeotto et al., “One-step design of a stable variant of the malaria invasion protein RH5 for use as a vaccine immunogen,” Proc. Natl. Acad. Sci., vol. 114, no. 5, pp. 998–1002, Jan. 2017, doi: 10.1073/pnas.1616903114.

[3] H. Lu et al., “Machine learning-aided engineering of hydrolases for PET depolymerization,” Nature, vol. 604, no. 7907, pp. 662–667, Apr. 2022, doi: 10.1038/s41586-022-04599-z.

[4] L. Scheibenreif, M. Littmann, C. Orengo, and B. Rost, “FunFam protein families improve residue level molecular function prediction,” BMC Bioinformatics, vol. 20, no. 1, p. 400, Dec. 2019, doi: 10.1186/s12859-019-2988-x.

[5] D. N. Woolfson, “A Brief History of De Novo Protein Design: Minimal, Rational, and Computational,” J. Mol. Biol., vol. 433, no. 20, p. 167160, Oct. 2021, doi: 10.1016/j.jmb.2021.167160.

[6] N. Metropolis and S. Ulam, “The monte carlo method,” J. Am. Stat. Assoc., vol. 44, no. 247, pp. 335–341, 1949, doi: 10.1080/01621459.1949.10483310.

[7] B. Kuhlman and P. Bradley, “Advances in protein structure prediction and design,” Nat. Rev. Mol. Cell Biol., vol. 20, no. 11, pp. 681–697, Nov. 2019, doi: 10.1038/s41580-019-0163-x.

[8] Ahdritz, Gustaf, Bouatta, Nazim, Kadyan, Sachin, Xia, Qinghui, Gerecke, William, and AlQuraishi, Mohammed, “OpenFold.” Zenodo, Nov. 12, 2021. doi: 10.5281/ZENODO.5709539.

[9] J. Jumper et al., “Highly accurate protein structure prediction with AlphaFold,” Nature, vol. 596, no. 7873, pp. 583–589, Aug. 2021, doi: 10.1038/s41586-021-03819-2.

[10] R. Wu et al., “High-resolution de novo structure prediction from primary sequence.” bioRxiv, Jul. 22, 2022. doi: 10.1101/2022.07.21.500999.

[11] I. R. Humphreys et al., “Computed structures of core eukaryotic protein complexes,” Science, vol. 374, no. 6573, p. eabm4805, Dec. 2021, doi: 10.1126/science.abm4805.

[12] M. Varadi et al., “AlphaFold Protein Structure Database: massively expanding the structural coverage of protein-sequence space with high-accuracy models,” Nucleic Acids Res., vol. 50, no. D1, pp. D439–D444, Jan. 2022, doi: 10.1093/nar/gkab1061.

[13] N. Anand et al., “Protein sequence design with a learned potential,” Nat. Commun., vol. 13, no. 1, p. 746, Feb. 2022, doi: 10.1038/s41467-022-28313-9.

[14] J. Dauparas et al., “Robust deep learning based protein sequence design using ProteinMPNN.” bioRxiv, Jun. 04, 2022. doi: 10.1101/2022.06.03.494563.

[15] A. Madani et al., “ProGen: Language Modeling for Protein Generation.” arXiv, Mar. 07, 2020. Accessed: Jul. 28, 2022. [Online]. Available: http://arxiv.org/abs/2004.03497

[16] N. Ferruz, S. Schmidt, and B. Höcker, “ProtGPT2 is a deep unsupervised language model for protein design,” Nat. Commun., vol. 13, no. 1, p. 4348, Jul. 2022, doi: 10.1038/s41467-022-32007-7.

[17] E. Nijkamp, J. Ruffolo, E. N. Weinstein, N. Naik, and A. Madani, “ProGen2: Exploring the Boundaries of Protein Language Models.” arXiv, Jun. 27, 2022. Accessed: Jul. 28, 2022. [Online]. Available: http://arxiv.org/abs/2206.13517

[18] L. Moffat, S. M. Kandathil, and D. T. Jones, “Design in the DARK: Learning Deep Generative Models for De Novo Protein Design.” bioRxiv, Jan. 28, 2022. doi: 10.1101/2022.01.27.478087.

[19] D. Lowe, “Why AlphaFold won’t revolutionise drug discovery,” Chemistry World, Aug. 05, 2022. https://www.chemistryworld.com/opinion/why-alphafold-wont-revolutionise-drug-discovery/4016051.article (accessed Aug. 07, 2022).

[20] K. K. Yang, Z. Wu, and F. H. Arnold, “Machine-learning-guided directed evolution for protein engineering,” Nat. Methods, vol. 16, no. 8, pp. 687–694, Aug. 2019, doi: 10.1038/s41592-019-0496-6.

[21] M. Defresne, S. Barbe, and T. Schiex, “Protein Design with Deep Learning,” Int. J. Mol. Sci., vol. 22, no. 21, p. 11741, Jan. 2021, doi: 10.3390/ijms222111741.

[22] A. Elnaggar et al., “ProtTrans: Towards Cracking the Language of Lifes Code Through Self-Supervised Deep Learning and High Performance Computing,” IEEE Trans. Pattern Anal. Mach. Intell., pp. 1–1, 2021, doi: 10.1109/TPAMI.2021.3095381.

[23] P. Gainza, H. M. Nisonoff, and B. R. Donald, “Algorithms for protein design,” Curr. Opin. Struct. Biol., vol. 39, pp. 16–26, Aug. 2016, doi: 10.1016/j.sbi.2016.03.006.

[24] F. Morcos et al., “Direct-coupling analysis of residue coevolution captures native contacts across many protein families,” Proc. Natl. Acad. Sci., vol. 108, no. 49, Dec. 2011, doi: 10.1073/pnas.1111471108.

[25] R. Das and D. Baker, “Macromolecular Modeling with Rosetta,” Annu. Rev. Biochem., vol. 77, no. 1, pp. 363–382, Jun. 2008, doi: 10.1146/annurev.biochem.77.062906.171838.

[26] P.-S. Huang, S. E. Boyken, and D. Baker, “The coming of age of de novo protein design,” Nature, vol. 537, no. 7620, pp. 320–327, Sep. 2016, doi: 10.1038/nature19946.

[27] C. P. Hill, D. H. Anderson, L. Wesson, W. F. DeGrado, and D. Eisenberg, “Crystal structure of alpha 1: implications for protein design,” Science, vol. 249, no. 4968, pp. 543–546, Aug. 1990, doi: 10.1126/science.2382133.

[28] B. Lovejoy, S. Choe, D. Cascio, D. K. McRorie, W. F. DeGrado, and D. Eisenberg, “Crystal structure of a synthetic triple-stranded alpha-helical bundle,” Science, vol. 259, no. 5099, pp. 1288–1293, Feb. 1993, doi: 10.1126/science.8446897.

[29] A. Courbet et al., “Computational design of mechanically coupled axle-rotor protein assemblies,” Science, vol. 376, no. 6591, pp. 383–390, Apr. 2022, doi: 10.1126/science.abm1183.

[30] A. W. Senior et al., “Improved protein structure prediction using potentials from deep learning,” Nature, vol. 577, no. 7792, pp. 706–710, Jan. 2020, doi: 10.1038/s41586-019-1923-7.

[31] A. Kryshtafovych, T. Schwede, M. Topf, K. Fidelis, and J. Moult, “Critical assessment of methods of protein structure prediction (CASP)—Round XIII,” Proteins Struct. Funct. Bioinforma., vol. 87, no. 12, pp. 1011–1020, 2019, doi: 10.1002/prot.25823.

[32] M. AlQuraishi, “A watershed moment for protein structure prediction,” Nature, vol. 577, no. 7792, pp. 627–628, Jan. 2020, doi: 10.1038/d41586-019-03951-0.

[33] “Method of the Year 2021: Protein structure prediction,” Nature. https://www.nature.com/collections/dfejabhghd (accessed Aug. 05, 2022).

[34] J. Ingraham, V. Garg, R. Barzilay, and T. Jaakkola, “Generative models for graph-based protein design,” in Advances in neural information processing systems, 2019, vol. 32. [Online]. Available: https://proceedings.neurips.cc/paper/2019/file/f3a4ff4839c56a5f460c88cce3666a2b-Paper.pdf

[35] J. Ingraham, A. Riesselman, C. Sander, and D. Marks, “Learning protein structure with a differentiable simulator,” 2019. [Online]. Available: https://openreview.net/forum?id=Byg3y3C9Km

[36] A. Rives et al., “Biological structure and function emerge from scaling unsupervised learning to 250 million protein sequences,” bioRxiv, p. 622803, May 2019, doi: 10.1101/622803.

[37] M. Heinzinger et al., “Modeling aspects of the language of life through transfer-learning protein sequences,” BMC Bioinformatics, vol. 20, no. 1, p. 723, 2019.

[38] R. Rao et al., “Evaluating Protein Transfer Learning with TAPE,” in Advances in Neural Information Processing Systems 32, 2019, pp. 9689–9701. Accessed: Mar. 21, 2020. [Online]. Available: http://papers.nips.cc/paper/9163-evaluating-protein-transfer-learning-with-tape.pdf

[39] J. Meier, R. Rao, R. Verkuil, J. Liu, T. Sercu, and A. Rives, “Language models enable zero-shot prediction of the effects of mutations on protein function,” in Advances in neural information processing systems, 2021, vol. 34, pp. 29287–29303. [Online]. Available: https://proceedings.neurips.cc/paper/2021/file/f51338d736f95dd42427296047067694-Paper.pdf

[40] Z. Lin et al., “Language models of protein sequences at the scale of evolution enable accurate structure prediction,” BioRxiv Prepr. Serv. Biol., 2022, doi: 10.1101/2022.07.20.500902.

[41] M. Bernhofer and B. Rost, “TMbed: transmembrane proteins predicted through language model embeddings,” BMC Bioinformatics, vol. 23, no. 1, p. 326, Aug. 2022, doi: 10.1186/s12859-022-04873-x.

[42] C. Marquet et al., “Embeddings from protein language models predict conservation and variant effects,” Hum. Genet., Dec. 2021, doi: 10.1007/s00439-021-02411-y.

[43] D. Ilzhoefer, M. Heinzinger, and B. Rost, “SETH predicts nuances of residue disorder from protein embeddings,” BioRxiv Prepr. Serv. Biol., 2022, doi: 10.1101/2022.06.23.497276.

[44] M. Littmann, M. Heinzinger, C. Dallago, T. Olenyi, and B. Rost, “Embeddings from deep learning transfer GO annotations beyond homology,” Sci. Rep., vol. 11, no. 1, pp. 1–14, 2021.

[45] M. Heinzinger, M. Littmann, I. Sillitoe, N. Bordin, C. Orengo, and B. Rost, “Contrastive learning on protein embeddings enlightens midnight zone,” NAR Genomics Bioinforma., vol. 4, no. 2, Jun. 2022, doi: 10.1093/nargab/lqac043.

[46] H. Stärk, C. Dallago, M. Heinzinger, and B. Rost, “Light attention predicts protein location from the language of life,” Bioinforma. Adv., vol. 1, no. 1, Nov. 2021, doi: 10.1093/bioadv/vbab035.

[47] M. Littmann, M. Heinzinger, C. Dallago, K. Weissenow, and B. Rost, “Protein embeddings and deep learning predict binding residues for various ligand classes,” Sci. Rep., vol. 11, no. 1, pp. 1–15, 2021.

[48] V. Thumuluri, J. J. Almagro Armenteros, A. R. Johansen, H. Nielsen, and O. Winther, “DeepLoc 2.0: multi-label subcellular localization prediction using protein language models,” Nucleic Acids Res., vol. 50, no. W1, pp. W228–W234, Apr. 2022, doi: 10.1093/nar/gkac278.

[49] M. H. Høie et al., “NetSurfP-3.0: accurate and fast prediction of protein structural features by protein language models and deep learning,” Nucleic Acids Res., vol. 50, no. W1, pp. W510–W515, Jun. 2022, doi: 10.1093/nar/gkac439.

[50] N. Ferruz and B. Höcker, “Controllable protein design with language models,” Nat. Mach. Intell., vol. 4, no. 6, pp. 521–532, Jun. 2022, doi: 10.1038/s42256-022-00499-z.

[51] H. Wang, S. Feng, S. Liu, and S. Ovchinnikov, “Disentanglement of Entropy and Coevolution using Spectral Regularization.” bioRxiv, p. 2022.03.04.483009, Mar. 07, 2022. doi: 10.1101/2022.03.04.483009.

[52] F. McGee et al., “The generative capacity of probabilistic protein sequence models,” Nat. Commun., vol. 12, no. 1, Art. no. 1, Nov. 2021, doi: 10.1038/s41467-021-26529-9.

[53] L. V. Castorina, R. Petrenas, K. Subr, and C. W. Wood, “PDBench: Evaluating Computational Methods for Protein Sequence Design.” arXiv, Sep. 28, 2021. doi: 10.48550/arXiv.2109.07925.

[54] Z. Li, Y. Yang, E. Faraggi, J. Zhan, and Y. Zhou, “Direct prediction of profiles of sequences compatible with a protein structure by neural networks with fragment-based local and energy-based nonlocal profiles,” Proteins, vol. 82, no. 10, pp. 2565–2573, Oct. 2014, doi: 10.1002/prot.24620.

[55] J. O’Connell et al., “SPIN2: Predicting sequence profiles from protein structures using deep neural networks,” Proteins Struct. Funct. Bioinforma., vol. 86, no. 6, pp. 629–633, Jun. 2018, doi: 10.1002/prot.25489.

[56] S. Chen et al., “To Improve Protein Sequence Profile Prediction through Image Captioning on Pairwise Residue Distance Map,” J. Chem. Inf. Model., vol. 60, no. 1, pp. 391–399, Jan. 2020, doi: 10.1021/acs.jcim.9b00438.

[57] A. Krizhevsky, I. Sutskever, and G. E. Hinton, “ImageNet Classification with Deep Convolutional Neural Networks,” in Advances in Neural Information Processing Systems, 2012, vol. 25. Accessed: Aug. 28, 2022. [Online]. Available: https://proceedings.neurips.cc/paper/2012/hash/c399862d3b9d6b76c8436e924a68c45b-Abstract.html

[58] Y. Zhang et al., “ProDCoNN: Protein design using a convolutional neural network,” Proteins Struct. Funct. Bioinforma., vol. 88, no. 7, pp. 819–829, 2020, doi: 10.1002/prot.25868.

[59] G. Huang, Z. Liu, L. Van Der Maaten, and K. Q. Weinberger, “Densely Connected Convolutional Networks,” in 2017 IEEE Conference on Computer Vision and Pattern Recognition (CVPR), Jul. 2017, pp. 2261–2269. doi: 10.1109/CVPR.2017.243.

[60] Y. Qi and J. Z. H. Zhang, “DenseCPD: Improving the Accuracy of Neural-Network-Based Computational Protein Sequence Design with DenseNet,” J. Chem. Inf. Model., vol. 60, no. 3, pp. 1245–1252, Mar. 2020, doi: 10.1021/acs.jcim.0c00043.

[61] A. V. Kulikova, D. J. Diaz, J. M. Loy, A. D. Ellington, and C. O. Wilke, “Learning the local landscape of protein structures with convolutional neural networks,” J. Biol. Phys., vol. 47, no. 4, pp. 435–454, Dec. 2021, doi: 10.1007/s10867-021-09593-6.

[62] L. V. Castorina, K. Subr, and C. W. Wood, “TIMED-Design: Efficient Protein Sequence Design with Deep Learning.” Zenodo, Aug. 16, 2022. doi: 10.5281/zenodo.6997495.

[63] A. J. Li, V. Sundar, G. Grigoryan, and A. E. Keating, “TERMinator: A Neural Framework for Structure-Based Protein Design using Tertiary Repeating Motifs.” arXiv, Apr. 27, 2022. doi: 10.48550/arXiv.2204.13048.

[64] A. J. Li, M. Lu, I. Desta, V. Sundar, G. Grigoryan, and A. E. Keating, “Neural Network-Derived Potts Models for Structure-Based Protein Design using Backbone Atomic Coordinates and Tertiary Motifs.” bioRxiv, p. 2022.08.02.501736, Aug. 18, 2022. doi: 10.1101/2022.08.02.501736.

[65] A. Strokach, D. Becerra, C. Corbi-Verge, A. Perez-Riba, and P. M. Kim, “Fast and Flexible Protein Design Using Deep Graph Neural Networks,” Cell Syst., vol. 11, no. 4, pp. 402-411.e4, Oct. 2020, doi: 10.1016/j.cels.2020.08.016.

[66] B. Jing, S. Eismann, P. Suriana, R. J. L. Townshend, and R. Dror, “Learning from Protein Structure with Geometric Vector Perceptrons.” arXiv, May 15, 2021. doi: 10.48550/arXiv.2009.01411.

[67] G. A. Orellana, J. Caceres-Delpiano, R. Ibañez, M. P. Dunne, and L. Alvarez, “Protein sequence sampling and prediction from structural data.” bioRxiv, p. 2021.09.06.459171, Nov. 22, 2021. doi: 10.1101/2021.09.06.459171.

[68] M. McPartlon, B. Lai, and J. Xu, “A Deep SE(3)-Equivariant Model for Learning Inverse Protein Folding.” bioRxiv, p. 2022.04.15.488492, Apr. 16, 2022. doi: 10.1101/2022.04.15.488492.

[69] Y. Cao, P. Das, V. Chenthamarakshan, P.-Y. Chen, I. Melnyk, and Y. Shen, “Fold2Seq: A Joint Sequence(1D)-Fold(3D) Embedding-based Generative Model for Protein Design.” arXiv, Jun. 24, 2021. doi: 10.48550/arXiv.2106.13058.

[70] C. Hsu et al., “Learning inverse folding from millions of predicted structures.” bioRxiv, p. 2022.04.10.487779, Apr. 10, 2022. doi: 10.1101/2022.04.10.487779.

[71] N. Anand and P. Huang, “Generative modeling for protein structures,” in Advances in Neural Information Processing Systems, 2018, vol. 31. Accessed: Aug. 08, 2022. [Online]. Available: https://proceedings.neurips.cc/paper/2018/hash/afa299a4d1d8c52e75dd8a24c3ce534f-Abstract.html

[72] N. Anand, R. Eguchi, and P.-S. Huang, “Fully differentiable full-atom protein backbone generation,” Jul. 2022, Accessed: Aug. 22, 2022. [Online]. Available: https://openreview.net/forum?id=SJxnVL8YOV

[73] R. F. Alford et al., “The Rosetta all-atom energy function for macromolecular modeling and design,” J. Chem. Theory Comput., vol. 13, no. 6, pp. 3031–3048, Jun. 2017, doi: 10.1021/acs.jctc.7b00125.

[74] R. R. Eguchi, C. A. Choe, and P.-S. Huang, “Ig-VAE: Generative modeling of protein structure by direct 3D coordinate generation,” PLOS Comput. Biol., vol. 18, no. 6, p. e1010271, Jun. 2022, doi: 10.1371/journal.pcbi.1010271.

[75] B. Lai, M. McPartlon, and J. Xu, “End-to-End deep structure generative model for protein design.” bioRxiv, p. 2022.07.09.499440, Jul. 11, 2022. doi: 10.1101/2022.07.09.499440.

[76] S. Sabban and M. Markovsky, “RamaNet: Computational de novo helical protein backbone design using a long short-term memory generative neural network.” bioRxiv, Sep. 01, 2020. doi: 10.1101/671552.

[77] X. Guo, Y. Du, S. Tadepalli, L. Zhao, and A. Shehu, “Generating tertiary protein structures via interpretable graph variational autoencoders,” Bioinforma. Adv., vol. 1, no. 1, p. vbab036, Jun. 2021, doi: 10.1093/bioadv/vbab036.

[78] B. Huang et al., “A backbone-centred energy function of neural networks for protein design,” Nature, vol. 602, no. 7897, pp. 523–528, Feb. 2022, doi: 10.1038/s41586-021-04383-5.

[79] Z. Harteveld et al., “Deep sharpening of topological features for de novo protein design,” presented at the ICLR2022 Machine Learning for Drug Discovery, May 2022. Accessed: Aug. 12, 2022. [Online]. Available: https://openreview.net/forum?id=DwN81YIXGQP

[80] D. Ofer, N. Brandes, and M. Linial, “The language of proteins: NLP, machine learning & protein sequences,” Comput. Struct. Biotechnol. J., vol. 19, pp. 1750–1758, Jan. 2021, doi: 10.1016/j.csbj.2021.03.022.

[81] D. Repecka et al., “Expanding functional protein sequence spaces using generative adversarial networks,” Nat. Mach. Intell., vol. 3, no. 4, pp. 324–333, Apr. 2021, doi: 10.1038/s42256-021-00310-5.

[82] A. Vaswani et al., “Attention Is All You Need.” arXiv, Dec. 05, 2017. doi: 10.48550/arXiv.1706.03762.

[83] The UniProt Consortium, “UniProt: the universal protein knowledgebase in 2021,” Nucleic Acids Res., 2021, doi: 10.1093/nar/gkaa1100.

[84] A. Madani et al., “Deep neural language modeling enables functional protein generation across families.” bioRxiv, Jul. 18, 2021. doi: 10.1101/2021.07.18.452833.

[85] “Better Language Models and Their Implications,” OpenAI, Feb. 14, 2019. https://openai.com/blog/better-language-models/ (accessed Aug. 20, 2022).

[86] D. Hesslow, N. Zanichelli, P. Notin, I. Poli, and D. Marks, “RITA: a Study on Scaling Up Generative Protein Sequence Models.” arXiv, Jul. 14, 2022. doi: 10.48550/arXiv.2205.05789.

[87] J. Frazer et al., “Disease variant prediction with deep generative models of evolutionary data,” Nature, vol. 599, no. 7883, pp. 91–95, Nov. 2021, doi: 10.1038/s41586-021-04043-8.

[88] I. Anishchenko et al., “De novo protein design by deep network hallucination,” Nature, vol. 600, no. 7889, pp. 547–552, Dec. 2021, doi: 10.1038/s41586-021-04184-w.

[89] C. Szegedy et al., “Going Deeper with Convolutions.” arXiv, Sep. 16, 2014. doi: 10.48550/arXiv.1409.4842.

[90] D. Tischer et al., “Design of proteins presenting discontinuous functional sites using deep learning.” bioRxiv, Nov. 29, 2020. doi: 10.1101/2020.11.29.402743.

[91] C. Norn et al., “Protein sequence design by conformational landscape optimization,” Proc. Natl. Acad. Sci., vol. 118, no. 11, p. e2017228118, Mar. 2021, doi: 10.1073/pnas.2017228118.

[92] J. Wang et al., “Scaffolding protein functional sites using deep learning,” Science, vol. 377, no. 6604, pp. 387–394, Jul. 2022, doi: 10.1126/science.abn2100.

[93] M. Baek et al., “Accurate prediction of protein structures and interactions using a three-track neural network,” Science, vol. 373, no. 6557, pp. 871–876, Aug. 2021, doi: 10.1126/science.abj8754.

[94] N. Anand and T. Achim, “Protein Structure and Sequence Generation with Equivariant Denoising Diffusion Probabilistic Models.” arXiv, May 26, 2022. doi: 10.48550/arXiv.2205.15019.

[95] J. Sohl-Dickstein, E. A. Weiss, N. Maheswaranathan, and S. Ganguli, “Deep Unsupervised Learning using Nonequilibrium Thermodynamics.” arXiv, Nov. 18, 2015. doi: 10.48550/arXiv.1503.03585.

[96] J. Ho, A. Jain, and P. Abbeel, “Denoising Diffusion Probabilistic Models.” arXiv, Dec. 16, 2020. doi: 10.48550/arXiv.2006.11239.

[97] Y. Song and S. Ermon, “Generative Modeling by Estimating Gradients of the Data Distribution.” arXiv, Oct. 10, 2020. doi: 10.48550/arXiv.1907.05600.

[98] A. Ramesh, P. Dhariwal, A. Nichol, C. Chu, and M. Chen, “Hierarchical Text-Conditional Image Generation with CLIP Latents.” arXiv, Apr. 12, 2022. Accessed: Aug. 28, 2022. [Online]. Available: http://arxiv.org/abs/2204.06125

[99] T. Olenyi et al., “LambdaPP: Fast and accessible protein-specific phenotype predictions.” bioRxiv, p. 2022.08.04.502750, Aug. 05, 2022. doi: 10.1101/2022.08.04.502750.

[100] M. Mirdita, K. Schütze, Y. Moriwaki, L. Heo, S. Ovchinnikov, and M. Steinegger, “ColabFold: making protein folding accessible to all,” Nat. Methods, vol. 19, no. 6, Art. no. 6, Jun. 2022, doi: 10.1038/s41592-022-01488-1.

[101] M. van Kempen et al., “Foldseek: fast and accurate protein structure search.” bioRxiv, p. 2022.02.07.479398, Jun. 24, 2022. doi: 10.1101/2022.02.07.479398.

[102] N. Gohil, G. Bhattacharjee, K. Khambhati, D. Braddick, and V. Singh, “Engineering Strategies in Microorganisms for the Enhanced Production of Squalene: Advances, Challenges and Opportunities,” Front. Bioeng. Biotechnol., vol. 7, 2019, Accessed: Aug. 15, 2022. [Online]. Available: https://www.frontiersin.org/articles/10.3389/fbioe.2019.00050

[103] S. El-Gebali et al., “The Pfam protein families database in 2019,” Nucleic Acids Res., vol. 47, no. D1, pp. D427–D432, Jan. 2019, doi: 10.1093/nar/gky995.

[104] C. Rios-Martinez, N. Bhattacharya, A. P. Amini, L. Crawford, and K. K. Yang, “Deep self-supervised learning for biosynthetic gene cluster detection and product classification.” bioRxiv, p. 2022.07.22.500861, Jul. 23, 2022. doi: 10.1101/2022.07.22.500861.

[105] D. J. Newman and G. M. Cragg, “Natural Products as Sources of New Drugs from 1981 to 2014,” J. Nat. Prod., vol. 79, no. 3, pp. 629–661, Mar. 2016, doi: 10.1021/acs.jnatprod.5b01055.

[106] S. L. Schreiber, “The Rise of Molecular Glues,” Cell, vol. 184, no. 1, pp. 3–9, Jan. 2021, doi: 10.1016/j.cell.2020.12.020.

[107] L. Yao, Y. Zheng, and Z. Zhu, “Jasmonate suppresses seedling soil emergence in Arabidopsis thaliana,” Plant Signal. Behav., vol. 12, no. 6, p. e1330239, Jun. 2017, doi: 10.1080/15592324.2017.1330239.

[108] Q. L. Sievers et al., “Defining the human C2H2 zinc finger degrome targeted by thalidomide analogs through CRBN,” Science, vol. 362, no. 6414, p. eaat0572, 2018, doi: 10.1126/science.aat0572.

[109] E. S. Fischer, E. Park, M. J. Eck, and N. H. Thomä, “SPLINTS: Small-molecule protein ligand interface stabilizers,” Curr. Opin. Struct. Biol., vol. 37, pp. 115–122, Apr. 2016, doi: 10.1016/j.sbi.2016.01.004.

[110] U. K. Shigdel et al., “Genomic discovery of an evolutionarily programmed modality for small-molecule targeting of an intractable protein surface,” Proc. Natl. Acad. Sci., vol. 117, no. 29, pp. 17195–17203, Jul. 2020, doi: 10.1073/pnas.2006560117.

[111] D. Bier et al., “The Molecular Tweezer CLR01 Stabilizes a Disordered Protein–Protein Interface,” J. Am. Chem. Soc., vol. 139, no. 45, pp. 16256–16263, Nov. 2017, doi: 10.1021/jacs.7b07939.

[112] J. Rudolph, J. Settleman, and S. Malek, “Emerging Trends in Cancer Drug Discovery-From Drugging the ‘Undruggable’ to Overcoming Resistance,” Cancer Discov., vol. 11, no. 4, pp. 815–821, Apr. 2021, doi: 10.1158/2159-8290.CD-21-0260.

[113] S. A. Kautsar et al., “MIBiG 2.0: a repository for biosynthetic gene clusters of known function,” Nucleic Acids Res., vol. 48, no. D1, pp. D454–D458, Jan. 2020, doi: 10.1093/nar/gkz882.

[114] M. Piotrowski, P. Morsomme, M. Boutry, and C. Oecking, “Complementation of the Saccharomyces cerevisiae plasma membrane H+-ATPase by a plant H+-ATPase generates a highly abundant fusicoccin binding site,” J. Biol. Chem., vol. 273, no. 45, pp. 30018–30023, Nov. 1998, doi: 10.1074/jbc.273.45.30018.

[115] T. Jahn et al., “The 14-3-3 protein interacts directly with the C-terminal region of the plant plasma membrane H(+)-ATPase,” Plant Cell, vol. 9, no. 10, pp. 1805–1814, Oct. 1997, doi: 10.1105/tpc.9.10.1805.

[116] M. Marra, L. Camoni, S. Visconti, A. Fiorillo, and A. Evidente, “The Surprising Story of Fusicoccin: A Wilt-Inducing Phytotoxin, a Tool in Plant Physiology and a 14-3-3-Targeted Drug,” Biomolecules, vol. 11, no. 9, p. 1393, Sep. 2021, doi: 10.3390/biom11091393.

[117] F. H. Arnold, “Design by Directed Evolution,” Acc. Chem. Res., vol. 31, no. 3, pp. 125–131, Mar. 1998, doi: 10.1021/ar960017f.

[118] A. C. Hunt et al., “Multivalent designed proteins protect against SARS-CoV-2 variants of concern.” bioRxiv, Jul. 07, 2021. doi: 10.1101/2021.07.07.451375.

[119] P. C. Cirino and F. H. Arnold, “Exploring the Diversity of Heme Enzymes through Directed Evolution,” in Directed Molecular Evolution of Proteins, S. Brakmann and K. Johnsson, Eds. Weinheim, FRG: Wiley-VCH Verlag GmbH & Co. KGaA, 2002, pp. 215–243. doi: 10.1002/3527600647.ch10.

[120] V. de Crécy-lagard et al., “A roadmap for the functional annotation of protein families: a community perspective,” Database, vol. 2022, p. baac062, Jan. 2022, doi: 10.1093/database/baac062.

[121] E. Check Hayden, “The automated lab,” Nature, vol. 516, no. 7529, Art. no. 7529, Dec. 2014, doi: 10.1038/516131a.

[122] M. Segal, “An operating system for the biology lab,” Nature, vol. 573, no. 7775, pp. S112–S113, Sep. 2019, doi: 10.1038/d41586-019-02875-z.

[123] C. Arnold, “Cloud labs: where robots do the research,” Nature, vol. 606, no. 7914, pp. 612–613, Jun. 2022, doi: 10.1038/d41586-022-01618-x.

[124] “NVIDIA Omniverse for Digital Twins,” NVIDIA. https://www.nvidia.com/en-us/omniverse/solutions/digital-twins/ (accessed Aug. 23, 2022).

[125] F. Tao and Q. Qi, “Make more digital twins,” Nature, vol. 573, no. 7775, pp. 490–491, Sep. 2019, doi: 10.1038/d41586-019-02849-1.

[126] A. El Saddik, “Digital Twins: The Convergence of Multimedia Technologies,” IEEE Multimed., vol. 25, no. 2, pp. 87–92, Apr. 2018, doi: 10.1109/MMUL.2018.023121167.

[127] C. Krittanawong, “The next step in deep learning-guided clinical trials,” Nat. Cardiovasc. Res., vol. 1, no. 4, Art. no. 4, Apr. 2022, doi: 10.1038/s44161-022-00044-6.

[128] N. Zhou et al., “The CAFA challenge reports improved protein function prediction and new functional annotations for hundreds of genes through experimental screens,” Genome Biol., vol. 20, no. 1, p. 244, Nov. 2019, doi: 10.1186/s13059-019-1835-8.

[129] The Critical Assessment of Genome Interpretation Consortium, “CAGI, the Critical Assessment of Genome Interpretation, establishes progress and prospects for computational genetic variant interpretation methods.” arXiv, May 12, 2022. Accessed: Aug. 28, 2022. [Online]. Available: http://arxiv.org/abs/2205.05897

[130] S. Petti and S. R. Eddy, “Constructing benchmark test sets for biological sequence analysis using independent set algorithms,” PLOS Comput. Biol., vol. 18, no. 3, p. e1009492, Mar. 2022, doi: 10.1371/journal.pcbi.1009492.

[131] L. S. Lorello, A. Galassi, and P. Torroni, “BANANA: a Benchmark for the Assessment of Neural Architectures for Nucleic Acids,” Oct. 2021, Accessed: Aug. 07, 2022. [Online]. Available: https://openreview.net/forum?id=Pobz_8y2Q2_

[132] C. Dallago et al., “FLIP: Benchmark tasks in fitness landscape inference for proteins,” presented at the Thirty-fifth Conference on Neural Information Processing Systems Datasets and Benchmarks Track (Round 2), Jan. 2022. Accessed: Aug. 07, 2022. [Online]. Available: https://openreview.net/forum?id=p2dMLEwL8tF

[133] P. Notin et al., “Tranception: Protein Fitness Prediction with Autoregressive Transformers and Inference-time Retrieval,” in Proceedings of the 39th International Conference on Machine Learning, Jun. 2022, pp. 16990–17017. Accessed: Aug. 05, 2022. [Online]. Available: https://proceedings.mlr.press/v162/notin22a.html

[134] Z. Zhang et al., “Protein Representation Learning by Geometric Structure Pretraining.” arXiv, May 23, 2022. Accessed: Jul. 28, 2022. [Online]. Available: http://arxiv.org/abs/2203.06125

