## Supplement for "From sequence to function through structure: deep learning for protein design"

### Supplement 1

380,000 sequences were generated using ProtGPT2 using default parameters: top_k: 950, top_p: 1, temperature: 1, and max_length < 150 on an NVIDIA Quadro RTX 8000 with 48GB vRAM. The process took approximately 3.5h. From these sequences, 180,000 were automatically discarded because they had been truncated during the process (presented with no end-of-sentence tokens). We further ordered them by perplexity, and took the best 100,000 scoring ones. Predictions of functional and structural properties ran on the same GPU in approximately 30 minutes. The U50 dataset was generated by sampling 100,000 sequences from the Uniref50 database. The random dataset was created by randomly shuffling the amino acids in U50 sequences. Prediction of properties for these two datasets ran on the same GPU in ~1.5h/set.

Supplementary material

**From sequence to function through structure: deep learning for protein design**

Noelia Ferruz, Michael Heinzinger, Mehmet Akdel, Alexander Goncearenco, Luca Naef, Christian Dallago

### Supplement 2

Protein sequence of generated ProtGPT2 protein:

```
> seq6621
MARSILVTGANRGLGRGFIRQYLEQPNLHVTACIRDSASPAALDLAARHPGRLVVVELDISDEASVAAARSVLSEHQ
ITHLDVVVANAGLHTAFADLFSFTVDSALKHIDVNTIGTLRLIQALRPLVEKSPQPRFASMSSGMGSVADFTATPTF
NTYGYSVSKASLNMLTAKFHKEESWLKSFAVHPGFVLTDMGGDKAKLTVEEGVVGMVRVIEQATKESTGEFLRYDGT
PLPW
```

Check it online at:

https://embed.predictprotein.org/#/MARSILVTGANRGLGRGFIRQYLEQPNLHVTACIRDSASPAALDLAARHPGRLVVVELDISDEASVAAARSVLSEHQITHLDVVVANAGLHTAFADLFSFTVDSALKHIDVNTIGTLRLIQALRPLVEKSPQPRFASMSSGMGSVADFTATPTFNTYGYSVSKASLNMLTAKFHKEESWLKSFAVHPGFVLTDMGGDKAKLTVEEGVVGMVRVIEQATKESTGEFLRYDGTPLPW

Supplementary material

**From sequence to function through structure: deep learning for protein design**

Noelia Ferruz, Michael Heinzinger, Mehmet Akdel, Alexander Goncearenco, Luca Naef, Christian Dallago

### Supplement 3

We underline the potential of natural product space by showing and annotating the chemical space between 48 published molecular glues obtained from iPPIDB and known natural products in bacteria and fungi [1] (**Fig. S1**). We also showcase four examples of glue molecules very similar to four existing natural products (**Fig. S1, panels B-E**). Rapamycin, a macrocycle and BGC product extracted from Streptomyces hygroscopicus, discovered serendipitously on Easter Island (Rapa Nui) and currently used to prevent organ rejection, was identified to inhibit the metabolic master regulator mTOR (mammalian target of rapamycin) with exquisite selectivity by gluing it to the peptidyl-prolyl isomerase FKBP12, thus preventing access to the active site [2]. A non-natural product derived example is the case of lenalidomide, used in the treatment of multiple myeloma and the fifth most selling drug in 2021 [3], which has recently been discovered to act by gluing zinc-finger proteins such as the transcription factor CK1α, a class of proteins considered highly difficult to drug, to Cereblon [4], [5]. Cereblon belongs to a class of protein called E3-ligases which mark proteins with ubiquitin leading to their degradation.

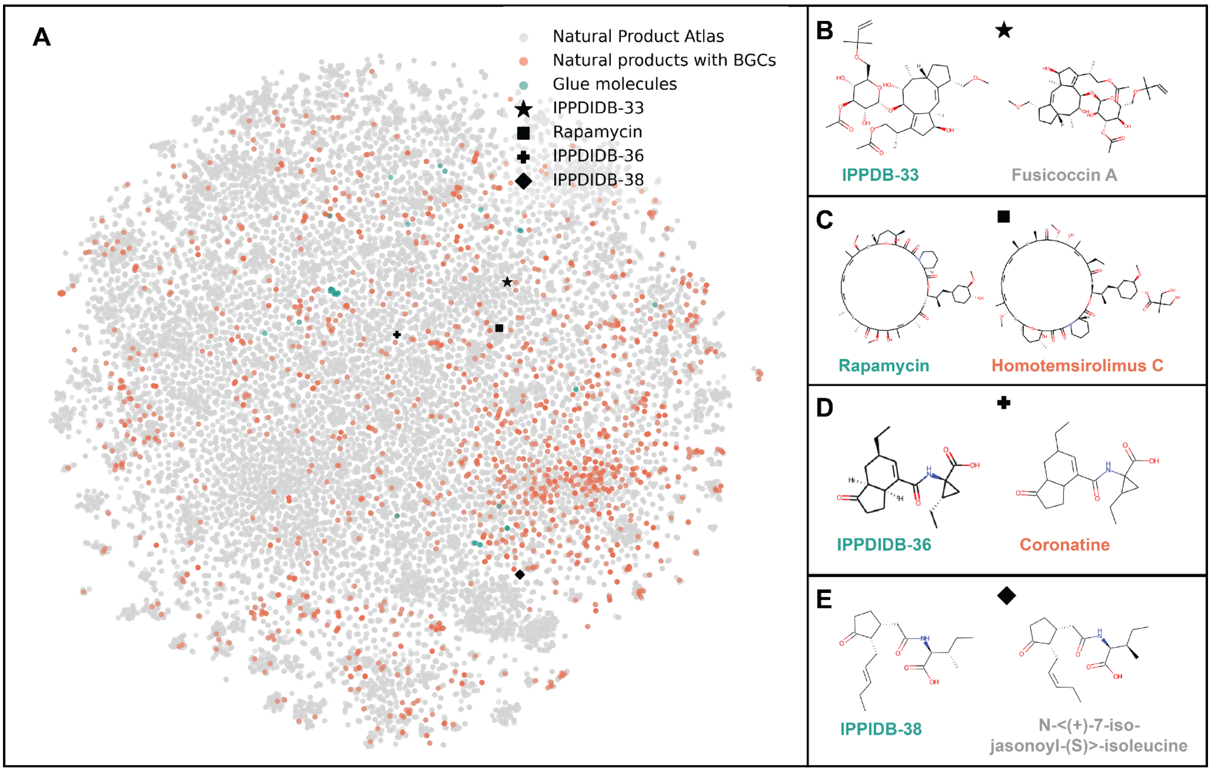


**Fig. S1**: a projection of t-SNE embeddings of Morgan chemical fingerprints for natural products from the *npatlas* [1] (gray), natural products from biosynthetic gene clusters (orange), and glue-like molecules (green) (panel A). Four examples of glue-like molecules are selected from panel A (marked ★, ■ ,✚ ,♦) cluster along with their closest natural products in panels B through E.

Supplementary material

**From sequence to function through structure: deep learning for protein design**

Noelia Ferruz, Michael Heinzinger, Mehmet Akdel, Alexander Goncearenco, Luca Naef, Christian Dallago

### References

[1] J. A. van Santen *et al.*, “The Natural Products Atlas 2.0: a database of microbially-derived natural products,” *Nucleic Acids Res.*, vol. 50, no. D1, pp. D1317–D1323, Jan. 2022, doi: 10.1093/nar/gkab941.

[2] J. Choi, J. Chen, S. L. Schreiber, and J. Clardy, “Structure of the FKBP12-Rapamycin Complex Interacting with Binding Domain of Human FRAP,” *Science*, vol. 273, no. 5272, pp. 239–242, Jul. 1996, doi: 10.1126/science.273.5272.239.

[3] B. Buntz, “50 of 2021’s best-selling pharmaceuticals,” *Drug Discovery and Development*, Mar. 29, 2022. https://www.drugdiscoverytrends.com/50-of-2021s-best-selling-pharmaceuticals/ (accessed Aug. 15, 2022).

[4] J. Krönke *et al.*, “Lenalidomide induces ubiquitination and degradation of CK1α in del(5q) MDS,” *Nature*, vol. 523, no. 7559, Art. no. 7559, Jul. 2015, doi: 10.1038/nature14610.

[5] Q. L. Sievers *et al.*, “Defining the human C2H2 zinc finger degrome targeted by thalidomide analogs through CRBN,” *Science*, vol. 362, no. 6414, p. eaat0572, 2018, doi: 10.1126/science.aat0572.
